# Uncovering zebrafish embryonic proteome dynamics across 16 time points during the first 24 hours of development

**DOI:** 10.64898/2026.03.24.713983

**Authors:** Fei Fang, William Poulos, Yifan Yue, Kun Li, Jose B Cibelli, Xiaowen Liu, Liangliang Sun

## Abstract

Defining how proteins change over developmental time is amenable to studies deciphering regulatory genetic networks in vertebrate development, biology, and pharmacology. In an approach toward such quantitative studies of dynamic network behavior, we produced an atlas using the mass spectrometry-based method to investigate protein expression changes across 16 time points from the zygote to the early pharyngula stage zebrafish embryos. We systematically summarize 8 clusters for interrogating changes in protein expression associated with the development of zebrafish embryos. Specifically, we identified a class of zinc finger-related transcription factors primarily located on the long arm of chromosome 4, which are highly expressed during zygotic genome activation. Furthermore, we highlight the power of this analysis to assign developmental stage-specific expression information to chromosomes and tissues. Time-resolved analyses reveal significant discordance between differential transcript and protein expression, whereas no time lag is observed for proteins involved in stable and fundamental biological processes, such as metabolism (e.g., Ppt2a and Gatm), cytoskeletal organization (e.g., Col18a1), and the translation machinery (e.g., Eif4enif1). This atlas offers high-resolution and in-depth molecular insights into zebrafish development, providing a resource for developmental biologists to generate hypotheses for functional analysis of proteins during early vertebrate embryogenesis.

**Highlights:**

- A global protein expression database with high time resolution is created for zebrafish embryos.
- Distinct patterns of protein expression correlate with biological processes.
- Transcription factors have a burst of expression from the gastrulation stage.
- Developmental stage-specific protein expressions were assigned to chromosomes and tissues.
- High-resolution embryonic transcriptome and proteome datasets were compared and connected.

## Introduction

Embryonic development is a dynamic period during which a fertilized egg undergoes cellular proliferation while simultaneously organizing into specific patterns and structures necessary to form tissues, organs, and the overall body of a new organism. Shortly after fertilization, animal embryos are transcriptionally quiescent and directed by maternal gene products to drive embryogenesis. During the maternal-to-zygotic transition (MZT) process, a large subset of maternally deposited gene products is cleared, and zygotic transcription is initiated, with developmental control handed over to those synthesized from the zygotic genome^1,2^. As development proceeds, global protein synthesis and differentiation are coordinated for dynamic morphological and functional changes.

During development, mechanical changes in cell and tissue shape drive morphogenesis, and gene expression changes regulate cell fate decisions and tissue patterning^3^. To understand development and create effective therapeutics, we need to grasp the dynamic, system-level interplay of genes and proteins across time, moving beyond individual gene or protein analysis.

The zebrafish, *Danio Rerio*, a unique vertebrate model system for biological research, has been mainly employed as an experimental model for embryogenesis, organogenesis, and general development in vertebrates^4^. It has many advantageous innate qualities, such as transparency, rapid embryonic development, and ease of manipulation and maintenance^5–7^. The high genomic conservation with humans it possesses, approximately 70% of the genes in the human protein-coding genome have *Danio Rerio* orthologs^8^, has also propelled the increasing characterization and analysis of zebrafish models to understand human diseases^9–13^.

Global gene expression profiling of the zebrafish using DNA microarrays offers insights into various embryonic and larval developmental stages. This has identified many genes that drive crucial steps of the differentiation process. For example, the maternally provided transcription factors Pou5f3, Sox19b, and Nanog were discovered to open up chromatin and prime genes for activity during zygotic genome activation in zebrafish with ribosome profiling, chromatin immunoprecipitation followed by parallel sequencing (ChIP-seq), and Assay for Transposase-Accessible Chromatin sequencing (ATAC-seq) strategies^14–16^. An mRNA expression time course of zebrafish development across 18 time points from 1 cell to 5 days post-fertilization (dpf) zebrafish embryo was produced, with temporal expression profiles of 23,642 genes characterized^17^. In addition, Madalena et al. provided a quantitative characterization of tRNA repertoires and their functional impact on decoding rates and codon-mediated mRNA decay during early zebrafish embryogenesis across 0 to 10 hours post-fertilization (hpf)^18^.

Compared with the ever-growing amount of data at the mRNA level, the proteome data involving the investigation of the physiological role of the complement system in zebrafish is virtually absent. However, accurate and complete pictures of physiological processes cannot be achieved through genomic information alone, as proteins are the executors of most of the biological functions. Thus, the study of the proteome, which provides temporal and spatial information of the translated genome, is imperative for understanding underlying biological mechanisms.

Currently, interest in mass spectrometry (MS)-based high-throughput proteomics technology, which could identify and quantify proteins and post-translational modifications, is growing rapidly used to investigate the proteins in zebrafish embryos^19–25^. The proteome of the embryo at several early developmental stages has been reported, providing important insights into the changes in the expression profiles of key proteins with an essential role in MZT^26,27^.

A time-resolved comprehensive analysis of relative protein expression levels is an important step toward understanding the regulatory processes governing embryonic development. Despite the crucial nature of this knowledge, we have yet to fully characterize the relative amount and dynamics of gene expression at the proteomic level due to the presence of a high abundance of yolk proteins in embryos. To the best of our knowledge, the only protein profiling across the whole zebrafish’s embryogenesis was conducted by Yan et al.^28^. With label-free quantitative proteomics, they identified 5961 proteins in zebrafish embryos across 10 time points from 4-cell stage to 5 dpf and revealed the expression dynamics of about 1000 proteins across the 10 time points. In their work, a 3-step deyolking and a single wash were performed, with 400-600 embryos used for each stage. However, such physical disruption procedures not only cause significant loss of material, which is not optimal for low-input proteomics, but also introduce technical variation in sample preparation that can impact downstream data interpretation. Our group recently performed quantitative proteomics of zebrafish embryos across four developmental stages (64-cell, 256-cell, dome, and 50% epiboly) during MZT without deyolking, revealing the expression dynamics of 5,000 proteins, representing one of the most systematic surveys of the proteome landscape of the zebrafish embryos during MZT^26^. Here, building upon our previous work, we measured protein expression throughout the zebrafish life cycle at 16 time points from 0 hpf to 24 hpf. To address the challenge of the large dynamic range of protein abundance, especially the large proportion of yolk protein in early embryos, we employed a high-resolution three-dimensional peptide fractionation before MS, including offline high pH reversed-phase liquid chromatography (RPLC) peptide fractionation, low-pH RPLC separation, and high-field asymmetric waveform ion mobility spectrometry (FAIMS) to characterize temporal expression profiles, which significantly increased the protein identification from early zebrafish embryos. The expression patterns of quantified 4418 proteins offer a comprehensive overview of protein-level changes associated with embryonic development. The proteins with significant abundance changes were clustered into eight groups, identifying the stage-specific proteins, especially the transcription factors, regulating embryo development in zebrafish. Our findings also revealed that maternal proteins showing significant abundance change before MBT were mainly located on chromosomes 3, 5, and 7. In addition, we found that mRNA and protein abundances are weakly correlated at the population gene level, while the metabolism, cytoskeletal organization, and translation machinery proteins show a strong correlation.

## Methods

### Material

All chemicals and reagents were of analytical grade and ordered from Sigma-Aldrich (St. Louis, MO) unless stated otherwise. Mammalian Cell-PE LB™ buffer for cell lysis was purchased from G-Biosciences (St. Louis, MO), while the complete mini protease inhibitor cocktail and phosphatase inhibitor were from Roche (Indianapolis, IN). A tandem mass tag (TMT) 16-plex reagent was ordered from Thermo Fisher Scientific (Rockford, IL). The MilliporeSigma™ Microcon-30 kDa Centrifugal Filter Unit with Ultracel-30 Membrane, as well as LC/MS grade water (H_2_O), acetonitrile (ACN), and formic acid (FA) were from Fisher Scientific (Pittsburgh, PA).

### Zebrafish maintenance and breeding

Following ZIRC/AALAC guidelines (https://www.aaalac.org/pub/?id=E9019693-90EC-FC4A-526E-E8236CC13B28), zebrafish are housed in a ZMod self-enclosed system. Water quality is monitored daily, including pH (maintained at 7.2-7.4), ammonia, and nitrite levels, to ensure optimal aquatic conditions and animal welfare. The water temperature was set at 28.5 °C, with a quarter of the water in the system changing daily. Each holding tank, made of polypropylene and with a 2-liter water capacity, can accommodate up to 12 adult fish. Almost every 4 weeks, fish are moved to a clean tank and kept under traditional conditions: 14h light/10h dark cycle. One day before the breeding day, we place a male and a female fish in the breeding tank, with a sieve for separation. On the breeding day, with the lights coming on, we removed the separator and allowed the fish to breed naturally. After natural breeding was observed, the fish were separated to manually collect the sperm and eggs for in vitro fertilization. Once the sperm and eggs were collected, the fish were placed back into tanks and rested for at least one month before the next round of breeding or experimental handling. The maintenance and breeding of zebrafish were performed by the Cibelli group in the Department of Animal Science at Michigan State University.

### Embryo collection

We collected 12 embryos at each of 16 different early stages (1-cell, 2-cell, 16-cell, 128-cell, 1k-cell, oblong, dome, 50% epiboly, shield, 75% epiboly, bud, 6-somite, 14-somite, 21-somite, 26-somite, and prim-5) with three independent biological replicates. The embryos were washed with cold phosphate-buffered saline buffer (Fisher Scientific, Pittsburgh, PA) twice. With the redundant liquid carefully removed, the embryos were immediately snap-frozen in liquid nitrogen and stored at -80 °C before use.

### Sample preparation for mass spectrometry analysis

For embryos at each stage, 120 µL of mammalian cell-PE LB^TM^ cell lysis buffer containing complete protease inhibitor and phosphatase inhibitor (pH 8.0) was added. To extract the proteins, the embryos were sonicated on ice using a Branson Sonifier 250 (VWR Scientific, Batavia, IL) for 2 min 6 times. With samples centrifuged at 16,000 g for 15 min at 4°C, the supernatants were collected to measure the protein concentration with the BCA assay (Thermo Fisher Scientific, Waltham, MA). Around 150 μg of total protein lysate from each sample was diluted to 0.8 μg/μL with lysis buffer and purified by acetone precipitation: 1 volume of protein sample was mixed with 4 volumes of pre-chilled (-20°C) acetone and the mixtures were kept at -20°C overnight. The tubes were then centrifuged at 14,000 g for 10 min, and the supernatants were discarded. The pellets were simply washed using 500 µL of cold acetone and re-centrifuged. The supernatants were discarded, and the protein pellets were placed in a chemical hood for 1∼2 min, leaving ∼5 µL of liquid. All the protein pellets were resuspended in 80 μL of 8 M urea in 50 mM NH_4_HCO_3_, and the protein concentration was measured with the BCA assay. For 50 μg of protein, denatured at 37°C for 1 h. Lysate was reduced by adding 1 μL of 0.4 M Dithiothreitol (DTT) and incubated at 56°C for 1 h, followed by alkylation by adding 1 μL of 1 M iodoacetamide (IAA) and incubating for 30 minutes in the dark. Then the mixture of each tube was loaded onto a 30-kDa centrifugal filter unit (250 µL/unit), followed by centrifugation at 14,000 g for 10 min. The proteins on the membrane were washed with 250 µL of 50 mM NH_4_HCO_3_ three times. Finally, 100 µL of 50 mM NH_4_HCO_3_ (pH 8.0) was loaded on each membrane, and 2 µL of trypsin solution (1 µg/µL) was added to each unit. The filter units were gently vortexed for 5 min to mix the trypsin and proteins. After that, the filter units were kept in a 37 °C water bath overnight for tryptic digestion. After digestion, the units were centrifuged at 14,000 g for 15 min, and the flow-through containing the peptides was collected. To increase peptide recovery from the membrane, the membrane was further washed with 50 µL of 50 mM NH_4_HCO_3_ twice. FA was used to acidify the protein digests to terminate digestion (1% FA final), followed by lyophilizing with a vacuum concentrator (Thermo Fisher Scientific). The digests were stored at -80 °C before use.

### TMT Isobaric Reagent labeling followed by high-pH RPLC fractionation

The peptides from embryos at the 16 embryonic stages were labeled with TMT16-plex labeling reagents, respectively, following the manufacturer’s protocol. Briefly, 20 µg of each lyophilized digest was dissolved in 20 µL of 200 mM HEPES buffer (pH 8.5). 8 µL of ACN was added to 50 µg of TMT reagent. The embryo digest sample from each developmental stage was transferred to the corresponding TMT channel vial and incubated for 1h at RT (**Fig. 1A**). 1 µL of 5% ethanolamine buffer was added to each vial, followed by incubation at room temperature for 15 min to block the residual TMT reagent. Then the digests were mixed and lyophilized to remove the organic solvent. When there was ∼10 µL of solution left, the lyophilization was stopped.

**Figure 1.**
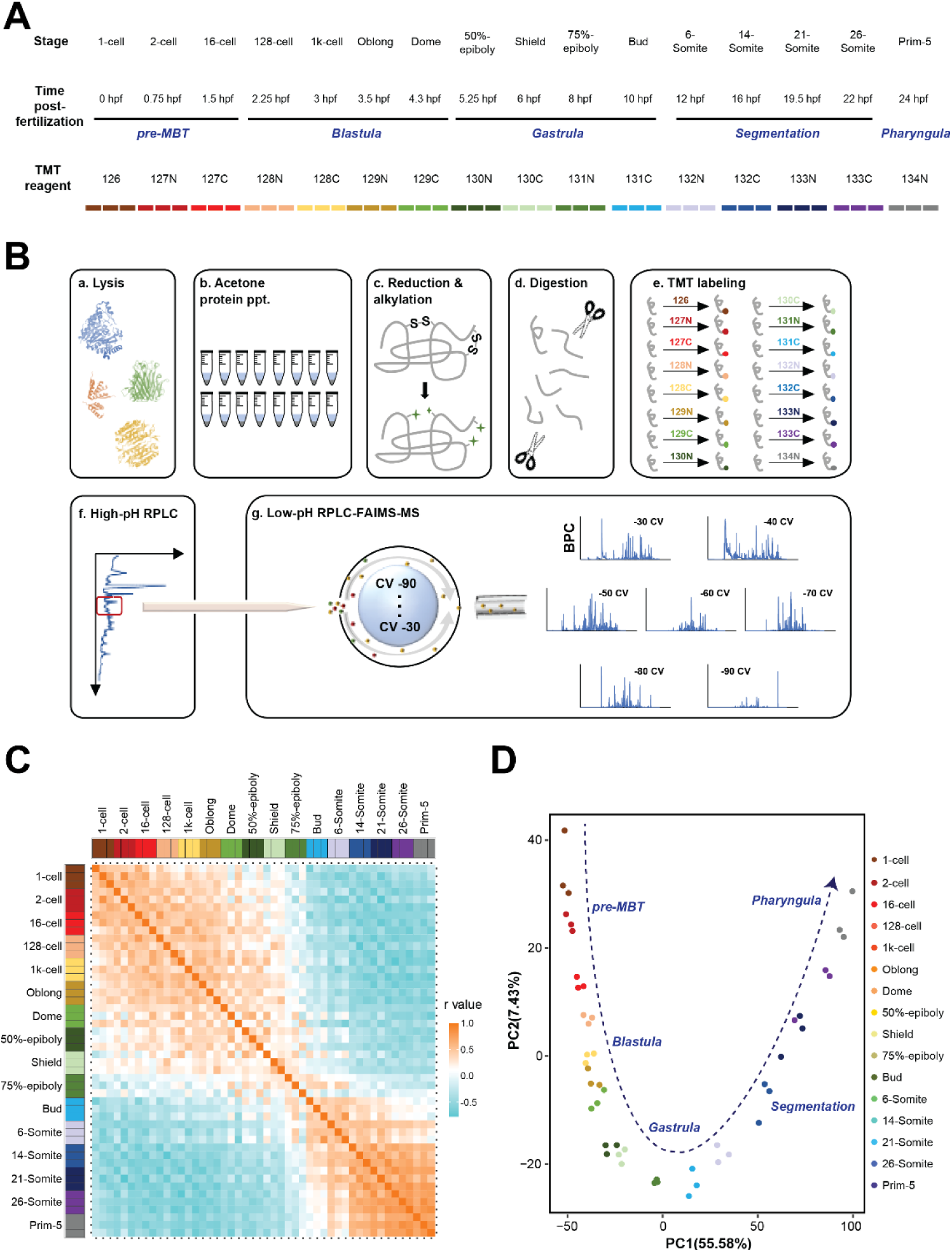
Summary of the quantified proteins from zebrafish embryos by RPLC-MS/MS. (A) Stages represented in this study. Also shown are the stage names, approximate hours post-fertilization (at 28.5°C) for each stage, five developmental categories, and different TMT tags used for each stage. The color scheme is used throughout the figures. (B) Workflow of the TMT-based quantitative experiment, encompassing protein extraction from the embryos at 16 different stages in biological triplicate (a), protein precipitation with acetone (b), protein reduction and alkylation (c), followed by tryptic digestion (d). The peptides labeled with 16 channels of TMT reagent (e) were mixed and pre-fractionated with offline high-pH RPLC (f), the fractions were further separated with online low-pH RPLC and FAIMS, followed by high-resolution MS analysis (g). (C) Sample correlation matrix of Spearman correlation coefficients of any two samples produced by comparing TMT report ion intensities (expression levels) across all proteins. (D) The 2D plot of Principal Component Analysis (PCA) shows the first two principal components (PCs). Samples from the same stage cluster together and there is smooth progression through developmental time (shown as a dashed arrow). Samples are colored as in (A) and are annotated with the stage categories. The amount of variance explained by each PC is indicated on each axis.

An Agilent Infinity II HPLC system was used for high-pH RPLC fractionation. A Zorbax 300Extend-C18 RP column (2.1 mm i.d. × 150 mm length, 3.5 µm particles, Agilent Technologies) was used for separation. Mobile phase A was 5 mM NH_4_HCO_3_ in water with pH 9, and mobile phase B was 5 mM NH_4_HCO_3_ in 80% ACN with pH 9. 200 µg TMT-labeled zebrafish embryo digest was injected into the RP column. The flow rate was 0.3 mL/min. The gradient was as follows: 0-10 min, 6% B; 10-15 min, 6-10% B; 15-75 min, 10-50% B; 75-77 min, 50-100% B; 77-87 min, 100% B; 87-90 min, 100-2% B; 90-100 min, 2% B. In total, 60 fractions were collected from 15 min to 75 min, one fraction per minute. Another fraction was collected from 75 min to 85 min. We named the fractions based on their retention times, ranging from 1 to 60. Then we combined fraction N, fraction N+20, and fraction N+40 to generate the first 20 fractions. The remaining fraction was named as fraction 21 (**Figure S1**). Those fractions were then lyophilized and stored at -80 °C for low-pH RPLC-FAIMS*-*MS/MS analysis.

### RPLC-FAIMS-MS/MS for the TMT-labeled sample

An EASY-RPLC™ 1200 system (Thermo Fisher Scientific) was used for RPLC separation. Each of the 20 fractions was dissolved in 28 µL of 0.1% (v/v) FA and 2% (v/v) ACN. 1 µL of the sample was injected and separated on a C18 separation column (75-µm i.d. × 50 cm, ReproSil-Pur 120 C18-AQ, 1.9 µm particles, 120 Å, Thermo Scientific) at a flow rate of 180 nL/min, held at 50 °C during separation by a column oven. Mobile phase A was H_2_O containing 2% (v/v) ACN and 0.1% (v/v) FA, while mobile phase B was H_2_O containing 80% (v/v) ACN and 0.1% (v/v) FA. For separation, a 70-minute gradient was used: 0-50 min, 15-45% B; 50-55 min, 45-80% B; 55-62 min, 80% B, 62-63 min, 80-2% B, and 63-70 min, 2% B.

The FAIMS device was placed between the nanoelectrospray source and the mass spectrometer. FAIMS separations were performed with the following settings: inner electrode temperature = 100 °C, outer electrode temperature = 100 °C, FAIMS carrier gas flow = 4.6 L/min, entrance plate voltage = 250 V, and CV settling time = 25 ms. The FAIMS carrier gas is N_2_ only, and the ion separation gap is 1.5 mm. The dispersion voltage (DV) circuitry was tuned using the auto-tune option. The option “Tune DV RF” was enabled throughout the RPLC-FAIMS-MS/MS analysis for the high and low electric field’s automated settings that create the DV waveform applied to the electrodes. Peptide mode CV ranging from -90 to -30 V or -100 to -40 V with 10 V steps was applied throughout the analysis.

An Orbitrap Exploris 480 mass spectrometer (Thermo Fisher Scientific) was used for the RPLC-FAIMS-MS/MS experiments. The ion transfer tube temperature was set at 320 °C, and the RF lens was set at 40%. The spray voltage was set to 2.0 kV. A Top15 DDA method was used. The mass resolution was set to 120,000 (at m/z 200) and 45,000 for full MS scans and MS/MS scans. For full MS scans and MS/MS scans, the target values were 3E6 and 1E5, and the maximum injection time was 100 ms and 160 ms, respectively. The scan range for MS scans was 350 to 1400 m/z. For MS/MS scans, the isolation window was 0.7 m/z. Fragmentation in the HCD cell was performed with a normalized collision energy of 32%. The fixed first mass was set to 100 m/z. Dynamic exclusion was applied, and it was set to 45 s. Ions with determined charge states between 2 to 5 were selected for fragmentation.

### Database search and data analysis

All raw files were analyzed by Proteome Discoverer 3.0 and MaxQuant 2.1.2.0 software with the Andromeda search engine^29^.

For Proteome Discoverer, the raw files were searched against the database of the *Danio rerio* proteome (ID: UP000000437, 46,848 entries, Oct 12, 2021) downloaded from UniProt (http://www.uniprot.org/). To calculate the false positive rate (FDR), inverse libraries were added to the database. The FDR for identification at the spectral, peptide, and protein levels was set at 1%. The enzymatic cleavage method was set at Trypsin (Full), the number of missed cut sites was set at 2, the minimum peptide length was set at 6 amino acid residues, the maximum number of peptide modifications was set at 3, the mass error tolerance of the primary parent ion was set at 20 ppm and the mass error tolerance of the secondary fragment ion was set at 0.02 Da. The dynamic modifications were deamidation (N/Q), oxidation (M), acetylation on protein N-terminus, TMTpro (K), and TMTpro (N-Term), and carbamidomethyl (C) were set as static modifications.

MaxQuant was applied for further quantitative analysis. TMTpro on lysine and N-terminus were selected as peptide labels. The *Danio rerio* proteome (ID: UP000000437, 46,848 entries, Oct 12, 2021) downloaded from UniProt (http://www.uniprot.org/) was used for database search. The peptide mass tolerances of the first search and main search were 20 and 4.5 ppm, respectively. The fragment ion mass tolerance was 20 ppm. Trypsin was selected as the protease. The dynamic modification was deamidation (N/Q), oxidation (M), and acetylation on protein N-terminus, and carbamidomethyl (C), TMTpro (K), and TMTpro (N-Term) were set as static modifications. The minimum length of a peptide was set to 7. Match between runs was enabled. The FDRs were 1% for both peptide and protein.

### Statistical analysis

For the time-series proteome experiment, the statistical analysis was performed using the statistical software R (R version 4.2). Firstly, the database search results obtained from MaxQuant were respectively subjected to Perseus software (v1.6.15.0)^30^. All entries marked by MaxQuant as a “Potential contaminant”, “Reverse”, or “Only identified by site” were deleted, followed by the removal of the protein containing intensity as zero. To address the heterogeneity of variance, the quantile normalization transformation was performed on the reporter ion intensities from the RPLC-FAIMS-MS/MS method with Normalyzer^31^. Then, the normalized read counts were applied for temporal expression analysis using the maSigPro^32^ with a degree=2. The proteins with an adjusted FDR-corrected p-value<0.05 were determined as proteins with statistically significant abundance changes. Finally, the MFuzz model was employed to cluster together significant proteins with similar expression patterns and visualized using the ClusterGVis package^33^.

For some statistical analysis and data visualization, RStudio, along with the R language (v.4.2.2), was employed. Principal-component analysis (PCA) was conducted using the R function ‘princomp’^34^. The visualization task of PCA and correlation analysis was conducted by the package ‘ggplot2’^35^, and the chromosomal loci of the differentially expressed proteins were prepared with the package ‘circlize’^36^. Gene ontology (GO) functional classification was performed with the web-based DAVID Bioinformatics Resources (https://david.ncifcrf.gov/)^37,38^ and REVIGO (http://revigo.irb.hr/)^39^ with the default settings. The enrichment tool was used to analyze clusters of genes to find shared common biological features and functions using a p-value cutoff of 0.05. STRING (v.12.0) (https://string-db.org/)^40^ is applied to obtain the protein-protein interaction network analyses. Network type is a full STRING network, and the edges indicate both physical and functional associations of proteins. The network edge suggests the confidence of protein-protein connections, and the line thickness indicates the strength of data support, with thicker lines as more confident connections. The minimum required interaction score is medium confidence (0.4). Others are default settings and visualized using the Cytoscape software (v.3.10.2)^41^.

To calculate the correlation coefficient between transcriptome and proteome datasets, the normalized protein/RNA expression relative to the 1-cell stage was averaged at each stage (protein, n=3, RNA, n=5), then the Pearson correlation coefficients were calculated using the R functions ‘cor.test’^42^. The genes with a correlation coefficient larger than 0.6 with a p-value less than 0.05 are considered as significant positive correlation, while correlation coefficients less than -0.6 with a p-value less than 0.05 are considered as significant negative correlation.

## Results and Discussion

### A high temporal resolution proteome profile of zebrafish embryonic development

In this study, we sampled a total of 16 developmental time points, including three points before the mid-blastula transition (MBT), four during the blastula period, four during the gastrula period, four during the segmentation period, and one during the early pharyngula period (**Figure 1A**). This gives detailed coverage of all the important developmental processes taking place during the first day of the embryo’s life. Total protein was extracted at 16 different developmental time points (see Methods for details), and for each point analysis, three independent biological replicates were analyzed.

MS-based proteomics analysis of eggs and early embryos is often challenging due to the high yolk content. For instance, in early zebrafish embryos (through gastrulation), yolk constitutes ∼90% of egg protein content, limiting the depth of proteomics analysis^43^. As shown in **Figure S2**, with 1D RPLC-MS/MS analysis of a zebrafish embryo proteome digest, we only obtained 300-400 proteins per sample due to the highly abundant yolk proteins. The distribution of expression levels (iBAQ, intensity-based absolute quantification) is initially unimodal, with a shift toward higher-abundance proteins, becoming more evenly distributed as the embryo develops.

The above data indicated that the yolk protein severely limited the depth of proteomics analyses. Researchers usually remove the yolk through centrifugation after lysis of eggs or embryos^24,26^, which leads to the loss of development-related proteins^44^. Recognizing the vital role of sample preparation in increasing proteome depth tailored for early embryos, we address the pressing need for an efficient and reproducible method by integrating two additional separation mechanisms, including high-pH RPLC and FAIMS, which separate ions in the liquid phase and gas phase, respectively, for the deep proteome profiling (**Figure 1B**). To improve the analysis throughput and keep quantification accuracy, TMT 16 pro reagents^45^ were used to quantify proteins across 16 distinct developmental time points in one experiment.

FAIMS separation, which is orthogonal to both RPLC and MS, is used as a means of online fractionation to improve the detection of peptides in complex samples^46^. FAIMS separates gas-phase ions based on their characteristic differences in mobility in high and low electric fields, which could remove interfering ion species and select peptide charge states optimal for identification by MS/MS, **Figure S3A**. As shown in **Figure S3B**, while 34% to 48% of peptides overlap between adjacent CVs, the overlap can be as small as about 10% for CVs that are far apart. The above results demonstrate that FAIMS is efficient in reducing the proteomic complexity of zebrafish embryos and increasing the proteome depth. Attributed to the orthogonality in the separation mechanism of different methods, ∼7,800 proteins were identified in one biological replicate from ∼50,000 peptides with Proteome Discoverer (**Figure S3C**), representing the deepest proteome coverage of zebrafish early-stage embryos so far. For more accurate and reliable quantification, MaxQuant was employed for dataset analysis with normalization. About 5000 proteins were quantified in each biological replicate, **Figure S4A**, among which 4,418 proteins were confidently quantified across three biological replicates (**Supplementary File S2**). A sample correlation matrix produced by comparing expression levels across all proteins between samples further demonstrates excellent agreement among biological replicates of the same embryonic stage and substantial proteome shift as the embryo develops from the 1-cell stage to the Prim-5 stage, **Figure 1C**.

Principal component analysis (PCA) further shows close clustering of biological replicates and a smooth transition from one stage to another through developmental time, **Figure 1D**. The normalized protein expression relative to the 1-cell stage was averaged across biological triplicates at each stage. The distributions of normalized protein expressions become wider gradually from the 2-cell stage to the Prim-5 stage, **Figure S4B**, suggesting the protein expression changes are more significant as the embryos grow compared to the 1-cell stage, which agrees well with the embryonic phenotype changes.

We further compared our proteomics results with published Western blot data for several proteins^47,48^, **Figure S4C**. Three proteins were selected. Nanog, which activates zygotic gene expression during the MZT^14^, was found to increase the expression from the 1-cell to the dome stage, followed by a decrease. Smarce1, which does not affect the initial steps of development but is critical for endocardial development, heart tube formation, and heart looping, and posteriorly at later stages^49^, increases its expression across the blastula to the segmentation period. In addition, there’s no significant change across the whole developmental stages for Erk1, a member of the mitogen-activated protein kinase proteins, which is involved in dorsal-ventral patterning and subsequent embryonic cell migration^50^ and could be used as a loading control^47^. Our quantitative proteomics data align well with the published Western blot data, indicating the high reliability of our high-temporal-resolution zebrafish proteome dynamics dataset.

### Protein abundance profiling

By ordering the proteins in our dataset according to their maximum stage of expression, the broad expression of all proteins across development was visualized. As shown in **Figure 2A**, zebrafish embryos at each developmental stage have distinct pools of highly expressed proteins, indicating the drastic changes in the embryonic proteome during early development.

**Figure 2.**
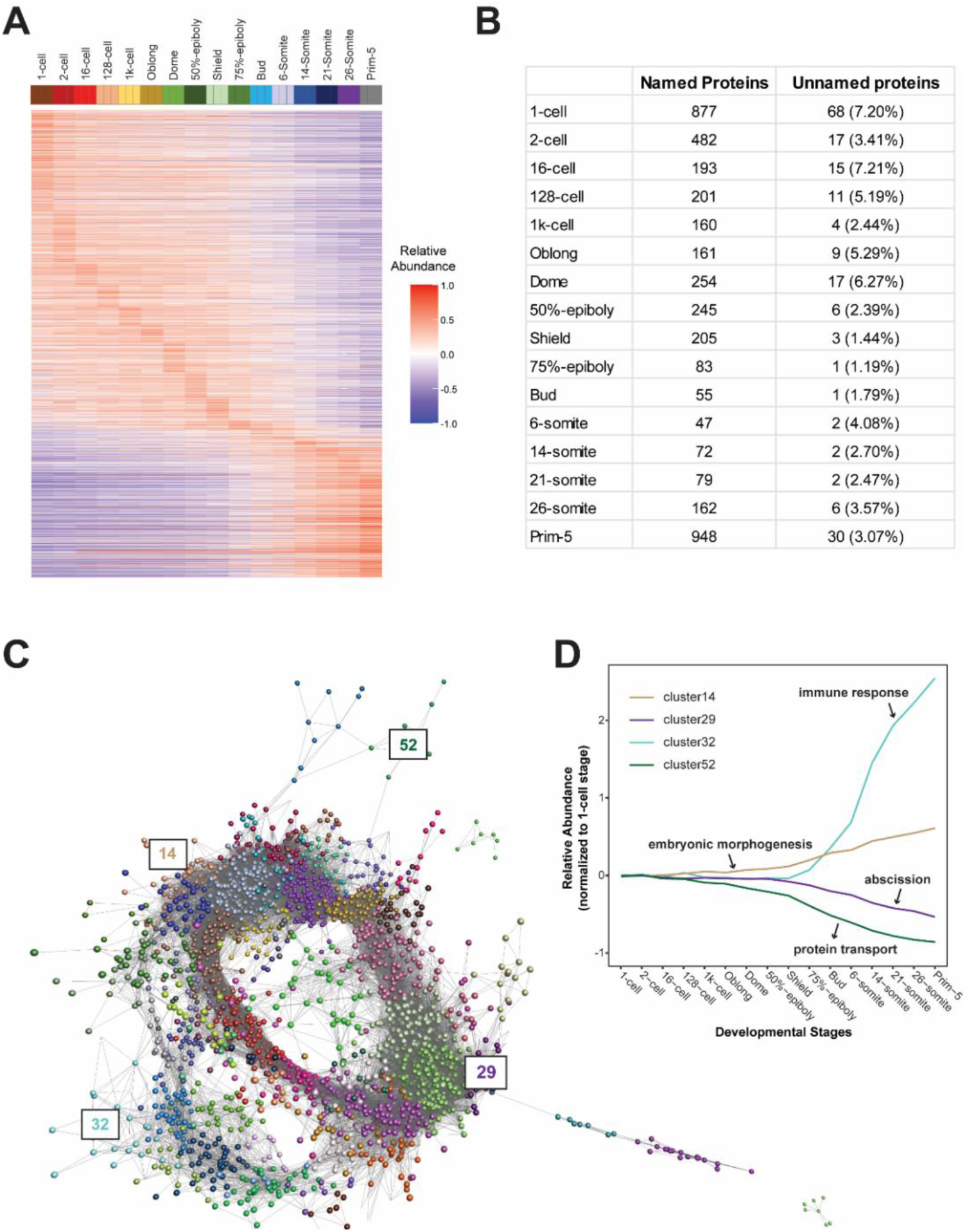
Clustering of expression patterns for all quantified proteins. (A) Heatmap of the expression profiles across the time course for all quantified proteins. The expression values are scaled to the maximum expression for that gene across the time course. Proteins are organized by the stage at which the maximum expression occurs and clustered within each stage. (B) Table of the numbers of named and unnamed proteins assigned to each stage. The unnamed proteins represent the proteins with undefined functions as well as lack orthologues to other species, and are exclusive to zebrafish, while the named proteins represent those with known functions and identified orthologues. (C) Network diagram of the clustering produced by MCL on a network graph generated by linking proteins with a Pearson correlation coefficient of >0.94 and removing unassigned proteins and clusters with five proteins or fewer. Different colors indicate distinct clusters, each consisting of proteins with similar expression trends. The numbers indicate the positions in the network diagram of the clusters shown in D. (D) Example clusters illustrating the progression of expression peaks through development displayed as the mean-centered and variance-scaled TPM for the genes in the clusters.

Among the 4418 quantified proteins in this dataset, 194 proteins have automatically generated names (often derived from clone IDs) such as zgc:*, loc*, and si:ch*. Interestingly, the proportion of unnamed proteins, which lack orthologues exclusive to zebrafish, decreases significantly after the onset of gastrulation (**Figure 2B**). Unlikely, the proportion of unnamed genes at dome and gastrula stages is nearly twice as high (32.9-42.2%) as at other stages (18.2-28.6%) in mRNA expression^17^, demonstrating the discordance between transcriptome and proteome datasets for zebrafish embryogenesis.

While broad, temporally organized grouping of protein expression is useful for high-level developmental views, smaller and more specific grouping provides deeper insights, especially for understanding the roles of uncharacterized proteins^17^. To get small clusters that are more likely to contain biologically related proteins, we used the BioLayout *Express*^3D^ software (http://www.biolayout.org/)^51^, which builds a network graph in which proteins are nodes with edges between those whose expression is correlated above a given threshold, to cluster and visualize the dataset (**Supplementary File S3**). The BioLayout *Express*^3D^ software discovered the proteins that are more highly connected with each other than with the rest of the graph. In total, 64 clusters were generated within the atlas, among which only 10 have more than 100 proteins (**Figure 2C**). Four example clusters are shown in **Figure 2D**. Clusters 29 and 52 contain proteins that are maternally supplied and then degraded. Cluster 14 has proteins involved in embryonic morphogenesis that accumulate across the developmental stages. The proteins in cluster 32, which accumulate until gastrulation, play a significant role in immune response, aligning well with the transcriptomic findings^17^.

BioLayout *Express*^3D^ software clustered all quantified proteins. To elucidate the key proteins that drive crucial steps of the differentiation process, we further performed the maSigPro to investigate the proteins differentially expressed during embryogenesis, with a total of 2,220 proteins discovered as differentially expressed proteins and clustered into 8 groups (**Figure 3 and Supplemental File S4**).

**Figure 3.**
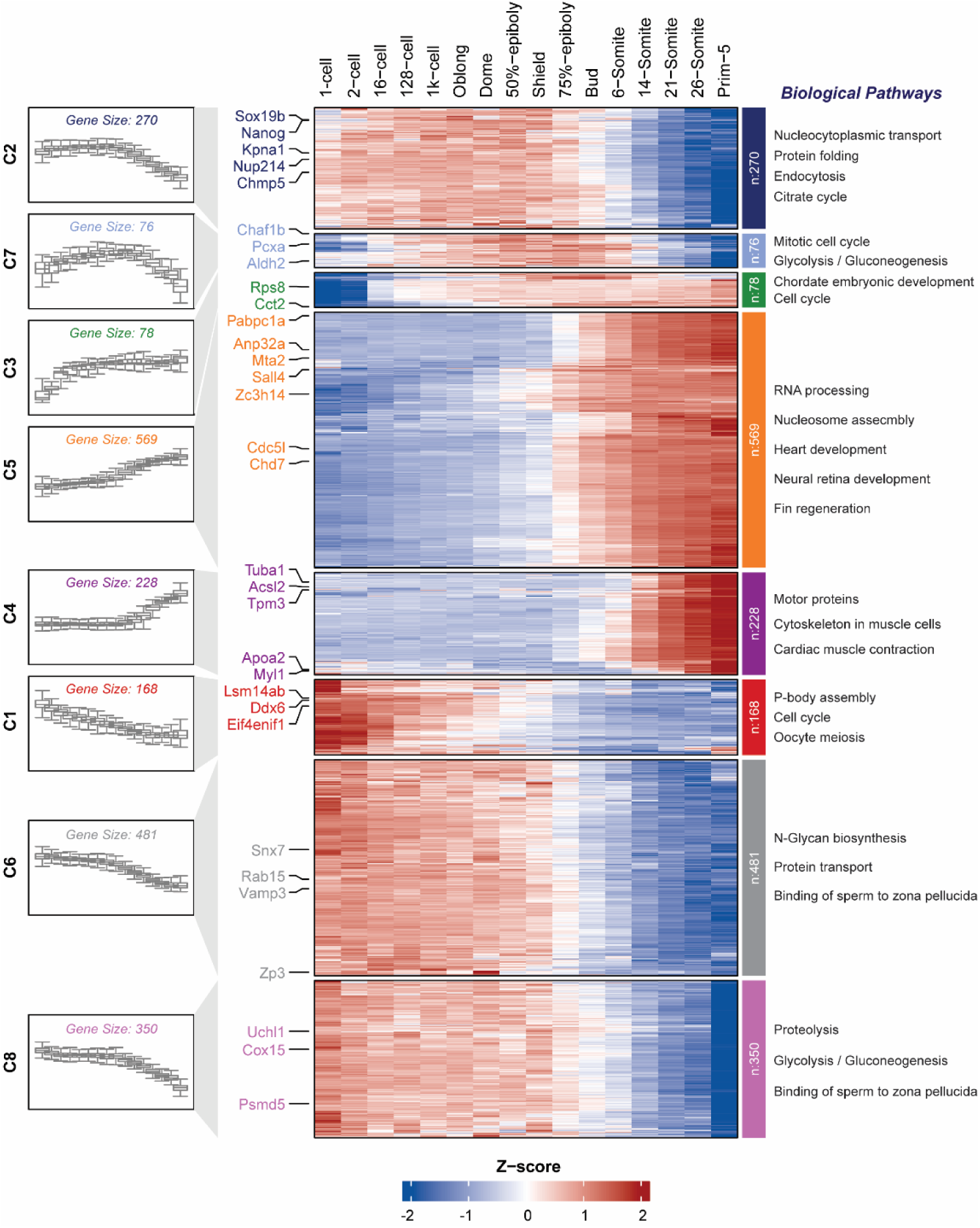
Cluster analysis of differentially expressed proteins. The heatmap in the middle shows distinct expression profiles of different clusters at each developmental stage. Fuzzy clustering of the expression data along the zebrafish development stages is shown at left. The functional enrichment data of the proteins in each cluster are shown to the right (p-value<0.05).

The proteins enriched in all clusters, apart from clusters 4 and 5, are those whose expression level significantly changes before MBT, which are most likely from maternal contribution. To confirm this point, we compared our proteomics data with the RNA-seq dataset by Bhat et al.^52^ and Lee et al.^14^, discovering that none of the proteins in these six clusters are strictly zygotic (**Supplemental File S4**), which are loaded into the egg during oogenesis, implement basic biosynthetic processes in the early embryo, direct the first mitotic divisions, and specify initial cell fate and patterning.

The 270 proteins enriched in cluster 2, whose expression level increases from pre-MBT and then gradually decreases from the gastrulation stage, include the most highly translated sequence-specific transcription factors (TFs) in the pre-MBT transcriptome, such as Nanog and Sox-19b, which open up chromatin and prime genes for activity during zygotic genome activation (ZGA) in zebrafish^14,16^. Another mechanism that accounts for the timing of ZGA onset is the nucleocytoplasmic ratio model, which hypothesizes that a ZGA repressor is present in the cytoplasm of the early embryo but is titrated by the increasing number of nuclei relative to the unchanging volume of cytoplasm^53^. We discovered that the proteins controlling nucleocytoplasmic transport, like nuclear pore complex protein (Nup214) and importin subunits (Kpna1, Kpna2, Kpna6, and Kpnb1), are highly enriched in cluster 2. In addition, abscission was recently reported to be a part of the MBT switch in early zebrafish embryogenesis^54^, while the components of the ESCRT machinery, Chmp2, Chmp4, Chmp5, and Vps4, which drive abscission, are found in cluster 2.

The 76 proteins in cluster 7 displayed a consistently ascending pattern followed by attenuation at the segmentation stage, playing essential roles in the mitotic cell cycle and glycolysis (**Supplemental File S5**). According to the RNA-seq datasets by Bhat et al.^52^, 40% of the proteins are gene products that are strictly maternally provided. Surprisingly, 50% of the proteins in cluster 7 are enzymes, such as strictly maternally provided oxidoreductase (Acat1, Decr1, Hadhab, and Acadm), aldehyde dehydrogenase (Aldh18a1, Aldh2, Aldh6a1, and Aldh9a1), and those involved in gluconeogenesis (Tpi1b and Pcxa) (**Table S1**). The aldehyde dehydrogenase superfamily represents a divergently related group of enzymes that metabolize a wide variety of endogenous and exogenous aldehydes^55^, among which Aldh2 was reported to have tissue-specific stem cell functions^56^. Most of the genes involved in gluconeogenesis are expressed in the extraembryonic yolk syncytial layer of the zebrafish embryo and contribute to the glucose supplementation to the embryos^57^, among which the deficiency of mitochondrial enzyme pyruvate carboxylase results in developmental delay^58^. In addition, the proteins involved in the mitotic cell cycle, such as chromatin assembly factor 1 (Chaf1), which functions as a histone chaperone that mediates the first step in nucleosome formation and whose loss results in delay and arrest of the cell cycle during organogenesis^59,60^, were enriched in cluster 7.

Cluster 3 contains 78 proteins, with a marked upregulation before the 16-cell stage, deciphering the early signaling events associated with MBT in zebrafish. In comparison with the RNA-seq datasets by Lee et al.^14^, all the proteins in cluster 3 are from maternal contribution, most of which are ribosomal proteins (41 out of 78), involved in protein synthesis, chordate embryonic development, and regulation of cell cycle (**Supplemental File S5**). In addition, the T-complex protein 1 subunits, Cct2 and Cct8, were found in cluster 3. The T-complex protein complex is involved in cell cycle regulation via substrate folding and protein interaction, transcription and translation initiation, as well as epigenetic modification^61^, indicating the potential role of the T-complex protein in regulating zygotic genome activation.

The proteins in clusters 5 and 4 were observed to significantly accumulate from the gastrulation and segmentation stages, respectively. We compared our proteomics data with the recent RNA-seq dataset by Bhat et al^52^, with pure zygotic proteins exclusively identified in these two clusters, aligning with the zygotic transcription required for gastrulation immediately after the MBT in zebrafish^62^. The zygotic proteins in these two clusters are enriched in tissue morphology, including heart, neural retina, and fin, coinciding with the fact that an organ- and tissue-level fate map is available for the onset of gastrulation^6^.

Interestingly, many transcription and translation regulators have been identified among cluster 5 proteins (Table S1), which regulate gene expression and protein synthesis. In addition, most of the proteins (316/519) in cluster 5 are in the nucleus (**Table S2**). The explosive increase in nuclear protein abundance is consistent with the initiation of ZGA and zygotic protein translation after the MBT, which requires numerous proteins involved in transcription and translation. We found that the proteins in this cluster are involved in polyadenylation, a process that occurs during MBT and is important for establishing normal zygotic transcription levels^63^. For example, zinc finger CCCH domain-containing protein 14 (Zc3h14), which is required for proper poly(A) tail length control, and poly(A) binding protein cytoplasmic 1 (Pabpc1), which plays a crucial role in the MBT by regulating the degradation of maternal mRNAs, are in cluster 5.

Post-translational processes were also reported to form part of the ZGA ‘timer’^1^. We found that phosphoproteins, which serve as key signaling molecules and regulators in various developmental processes such as cell movement, differentiation, and tissue formation, are significantly enriched in cluster 5. Among them, we identified acidic leucine-rich nuclear phosphoprotein 32 family members (Anp32a and Anp32e), which have been reported to influence stress-induced epigenetic regulation of gene expression and negatively influence cell cycle progression and proliferation^64^, as well as metastatic-associated proteins (Mta2 and Mta3), components of nucleosome remodeling and deacetylase (NuRD) complexes, function as an essential factor in pre-MBT *Xenopus* embryos^65^.

Among the 228 proteins enriched in cluster 4, motor proteins such as tubulin (Tuba1, Tubb2, Tuba8l2, and Tuba8l5), myosin (Myl1, Myl6, Myhc3, and Smyhc1) and tropomyosin (Tpm2, Tpm3 and Tpm4), which play a crucial role in regulating the dynamics of the actin cytoskeleton, particularly during cell migration, shape changes, and muscle development^66^, were significantly enriched. In addition, the proteins involved in peroxisome proliferator-activated receptor (PPAR) signaling, including apolipoprotein, long-chain-fatty-acid-CoA ligase, play a crucial role in systemic cell metabolism, energy homeostasis, and immune response inhibition.

The MZT is not only marked by the initiation of zygotic gene expression but also by the degradation of a subset of maternally provided mRNAs and proteins. Cluster 1 contains 168 proteins displaying a steady decline throughout embryogenesis, among which 109 proteins matched the SLAM-seq^52^. All proteins are from maternal contributions, and 44% of them are gene products that are strictly maternally provided. Among the transcripts expressed maternally provided and re-expressed in the zygote, 33 showed decreasing gene expression. The other proteins in cluster 1, such as RNA helicase Ddx6 and Ddx61, as well as mitotic checkpoint protein Bub3, are involved in P-body assembly, cell cycle, and oocyte meiosis, which agrees well with the transcriptome result^17^ and our previous proteomics data^27^. As many as 32% of the proteins in cluster 1 are unnamed proteins, whose functions might be predicted with our dataset in combination with protein function prediction methods such as DeepGO-SE, a method that predicts protein functions from protein sequences^67^.

The protein expression patterns in clusters 6 and 8 are very similar, with marked downregulation from the Bud stage, except that the proteins in cluster 8 decreased dramatically at the Prim-5 stage. Among the proteins in these two clusters, we identified zona pellucida sperm-binding proteins, which are important for inducing the acrosome reaction in sperm cells at the beginning of fertilization^68^. In addition, the proteins involved in proteolysis, such as proteasome subunits and ubiquitin carboxyl-terminal hydrolase, and complement components, a group of proteins that play a crucial role in the immune system, which are involved in the formation of unstable protease complexes^69^, are highly enriched in clusters 6 and 8.

In comparison with other clusters, proteins associated with the endomembrane system, including the endoplasmic reticulum, Golgi apparatus, endosomes, and lysosomes, were prominently enriched in cluster 6 (**Table S2**). The endomembrane system, such as the adaptor protein complex, sorting nexins, and vesicle-associated membrane proteins, plays a crucial role in the establishment of the three germ layers (ectoderm, mesoderm, and endoderm) by directing the movement and differentiation of cells through protein signaling. That might be because, during early cleavage divisions, the endomembrane system rapidly produces proteins and lipids needed for cell division and the formation of the blastocyst. Coincidentally, the membrane trafficking regulators, such as Rab GTPases (Rab1, Rab5, Rab6, Rab11, Rab13, Rab15, Rab18, and Rab33), which regulate molecule and protein transport between intracellular compartments and the extracellular milieu, were mostly enriched in cluster 6. The importance of Rab GTPase during development in vivo was reported on various metazoan model organisms, such as mice and C. elegans^70^. However, only Rab5 was reported to contribute to zebrafish embryo formation, such as the dorsal organizer^71^, and we found very few studies related to the roles of other Rab proteins during zebrafish embryonic development^72^.

Cluster 8 contains 350 proteins, mostly involved in metabolic pathways, and 72 of them are mitochondrial proteins (**Table S2**). Mitochondrial enzymes such as ATP synthase, which support the high energy demands required for cellular processes, such as differentiation, organogenesis, and rapid cell proliferation, decrease their expression at the prim-5 stage (**Table S1**).

Overall, we have produced a high-quality zebrafish embryonic proteome dynamics database. It represents the first detailed temporal protein expression reference in zebrafish, enhancing our understanding of the fundamental biological processes and events during early vertebrate development. The protein cluster data accurately reflect the main activities in embryonic development during pre-MBT, blastula, gastrula, segmentation, and early pharyngula periods. The proteome dynamics data also enable the visualization of expression changes in biological process-specific proteins across a broad spectrum of early developmental periods with high temporal resolution. The proteome resource offers developmental biologists pools of candidate proteins during early vertebrate development for studying the molecular mechanisms of specific biological questions. A web-based server has been created for the proteome dynamics resource generated in this study to facilitate the broad use of the dataset, and the server can be accessed through this link, https://www.toppic.org/software/zebrafishdb/zf_protein.html.

### A burst of transcription factor expression during MZT

One important model explaining the ZGA of zebrafish embryos is the accumulation of some transcriptional activators [e.g., transcription factors (TFs)] before the onset of ZGA^14^. The TFs are remarkably enriched in cluster 5, **Figure 4A**, demonstrating a burst of expression of TFs from the gastrulation stage. The zinc finger-related TFs represent the highest proportion (**Figure 4B**), among which 23% are located on chromosome 4. This finding is consistent with the transcriptome result that the zinc finger domain-containing genes, located on the long arm of chromosome 4, are expressed in a sharp peak during ZGA^17^.

**Figure 4.**
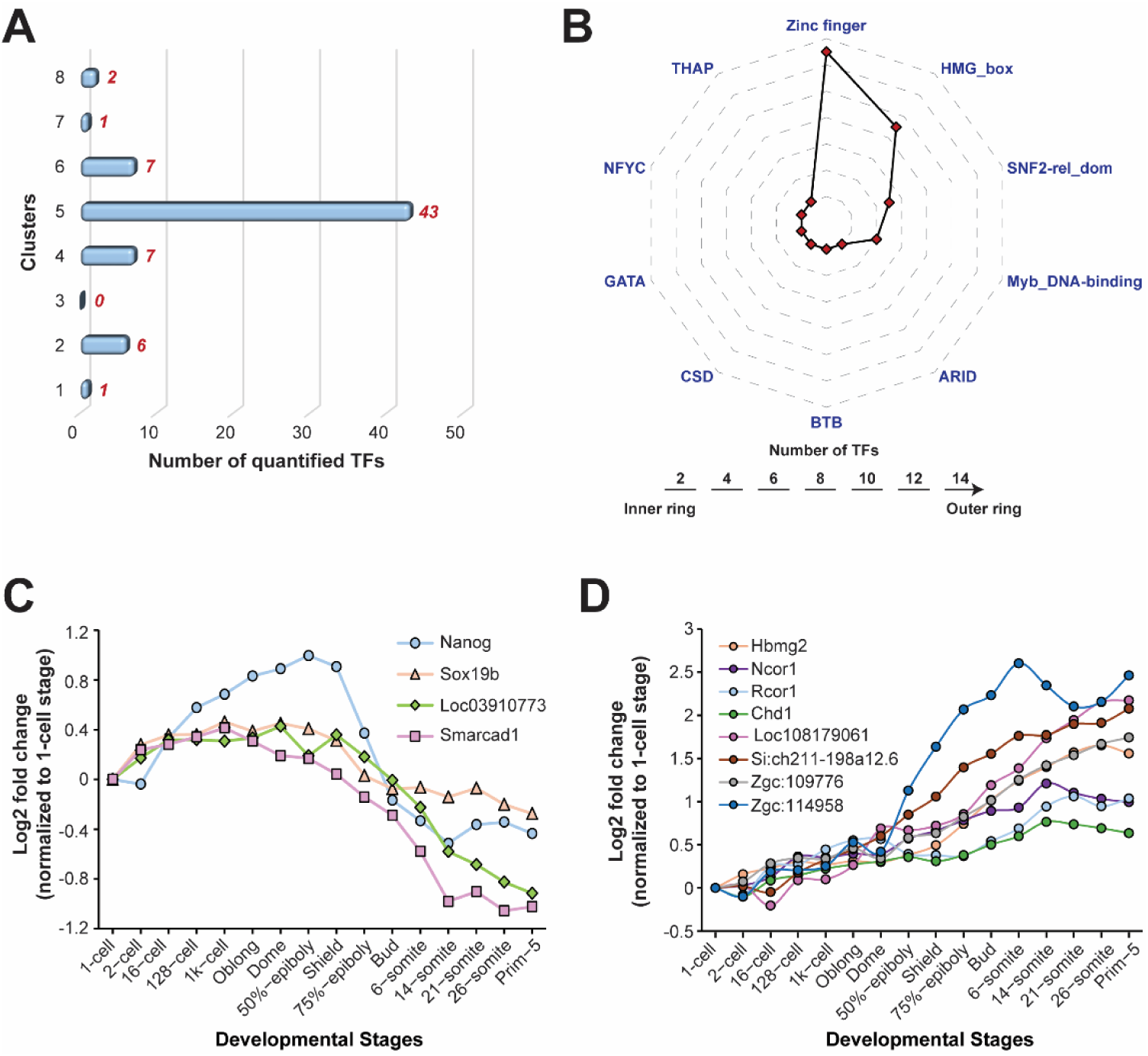
Expression level changes of the quantified transcription factors (TFs). (A) The quantified TFs and those with a significant change in expression level across development stages were classified into families according to their DNA-binding domain composition. Families with fewer than three members were excluded. (B) DNA-binding domain distribution of the TFs upregulated during the MZT. Expression profiles of TFs that show significant up-regulation before (C) and after (D) the onset of ZGA.

As shown in **Figure 4C**, four TFs exhibit a significant upregulation before the 16-cell stage, which may control the minor wave of ZGA before MBT. The expression patterns of two TFs, Nanog and Sox19b, which were reported to regulate ZGA in zebrafish^14,16^, agree well with the Western blot^47^ and RT-PCR result^73^. Although the function of Smarcad1 on zebrafish embryogenesis is unknown, the human SMARCAD1 is reported to be bound by the core pluripotency transcription factors NANOG, OCT4, and SOX2, with its increased expression correlating with naïve pluripotency and down-regulation upon differentiation^74–76^, indicating the potential role of zebrafish Smarcad1 in regulating ZGA.

Another eight TFs displayed a continuous increase, with a remarkable rise starting from the oblong stage, **Figure 4D**. Among them, Hmgb2, which is abundantly expressed during embryonic development^77^, shows a steady increase in protein expression levels during zebrafish gastrulation and early somitogenesis. With mouse HMGB2 acting upstream of OCT4/SOX2 signaling to control embryonic stem cell pluripotency^78^, the zebrafish homolog might share a similar function. We also discovered two corepressors, Ncor1 and Rcor1, which repress transcription by recruiting histone deacetylase^79,80^ and exhibit a continuous enhancement in protein expression before the segmentation stage. Ncor1 was shown to be crucial for the early patterning of the anterior-posterior axis of the zebrafish^81^. Maternal Rcor1 is essential for the repression of target genes during blastula stages, and Rcor1 is required to repress gene expression in mesodermal derivatives, including muscle and notochord, as well as within the nervous^82^. DNA helicase Chd1, which shows the same protein expression trend as Ncor1 and Rcor1, was reported to be associated with Ncor1 and histone deacetylase^80^. These transcriptional repressors might play a crucial role in the ZGA by acting as transcriptional co-repressors, helping to repress key genes that are necessary for the transition from early embryonic cleavage divisions to the onset of gastrulation, essentially acting as a “brake” on the rapid cell proliferation during early development and allowing for proper differentiation to begin.

Additionally, there are five other oocyte zinc finger TFs (i.e., names starting with “loc”, “Si”, or “zgc”) with unknown functions displaying an increase in expression during early embryogenesis (**Figures 4C** and **4D**), which might play essential roles in regulating early embryogenesis.

### Chromosome and tissue-related protein expression dynamics

We further mapped the differentially expressed proteins (DEPs) in each cluster onto each chromosome, **Figure 5**, aiming to observe chromosome-specific protein expression dynamics and understand the functional contributions of protein-coding genes on each chromosome to the zebrafish early embryonic development. The number of DEPs from each chromosome varied substantially, **Figure 5**. For example, chromosomes 1, 2, 3, 5, 7, 19, 20, and 21 have a relatively high number of DEPs, ranging from 97 to 138. Others have 59-92 DEPs. Chromosome 5 has the highest number of DEPs (138), and Chromosome 4 has the lowest number of DEPs (59). The data suggest that some chromosomes are more active than others during early embryogenesis.

**Figure 5.**
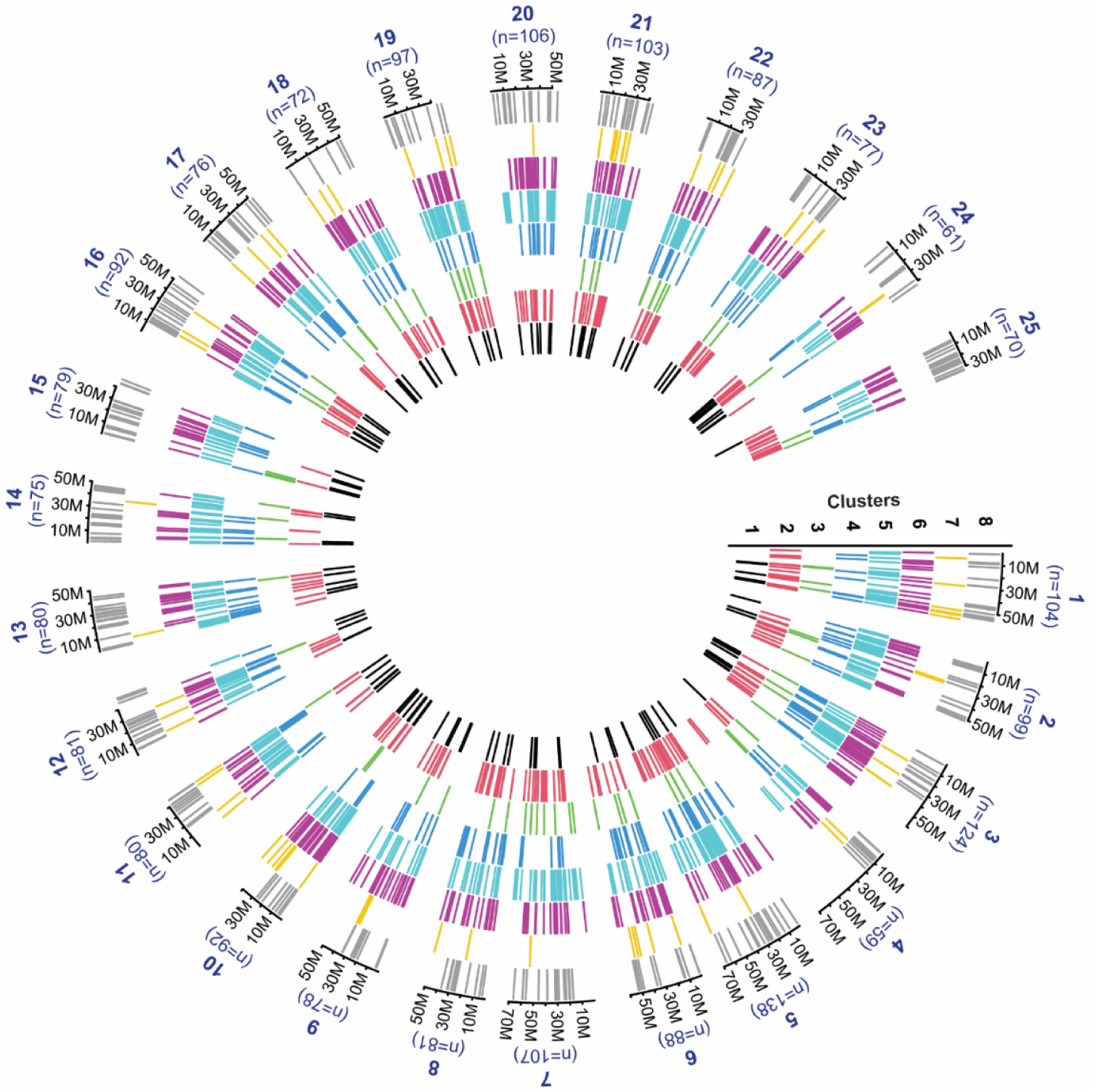
Chromosomal loci of the differentially expressed proteins (DEPs) from eight enriched protein expression clusters. Each bar represents each DEP, and different colors represent the DEP in different clusters. The protein cluster information is from Figure 3. The chromosome number and the total number of DEPs from each chromosome are labeled.

DEPs in each of the eight clusters of expression dynamics (**Figure 3**) do not distribute uniformly across the 25 pairs of chromosomes, **Figures 5** and **S5**. For example, for cluster 1 proteins, which are gradually degraded during development, chromosomes 3 and 24 contributed more than other chromosomes. For cluster 2 proteins, which are accumulated during the cleavage and blastula periods and then start to be degraded in the gastrula period, chromosome 5 contributes more than the others. For cluster 3 proteins, which gradually accumulated from the beginning of the development, chromosomes 7, 19, and 22 are the main contributors. Clusters 4 and 5 represent DEPs that accumulate starting from late or early gastrulation, respectively, and they exhibit similar distributions across chromosomes. Chromosome 5 contributes more than the others to clusters 4 and 5 proteins. Clusters 6 and 8 contain proteins that are both downregulated starting from the gastrula period and showing different chromosome distributions. Chromosomes 3, 10, and 20 contribute more than others to cluster 6 proteins; Chromosomes 5, 7, and 12-16 make more contributions to cluster 8 proteins. The results suggest that proteins from each chromosome have distinct functions in modulating early embryogenesis. This conclusion is further confirmed by the gene ontology (GO) analysis of DEPs from each chromosome. **Figure S6** shows five examples of chromosome data. Chromosome 5 DEPs are involved in ribosome biogenesis, cellular response to glucocorticoid stimulus, and mRNA processing, **Figure S6A**. Chromosome 7 DEPs play important roles in nucleosome assembly (histone H1 and H4), mRNA splicing (RNA binding proteins), and translation (ribosomal proteins), **Figure S6B**. Chromosome 3 DEPs are involved in mRNA splicing, nervous system development, and heart development, **Figure S6C**. The proteins encoded by genes on chromosome 22 primarily participate in cellular responses to estrogen stimuli and rRNA processing, **Figure S6D**. The genes on chromosome 24, including Nanog and Chmp4c, play important roles in reticulophagy, protein transport, and abscission, **Figure S6E**.

During organogenesis, proteins are up- and down-regulated in specific tissues. Although protein expression patterns of adult zebrafish organs were presented in several studies^83–85^, how the time-course proteomic states correlate with each other in the background of tissue localization in a complex developing zebrafish embryo remains elusive.

To construct the developmental trajectory, we integrated our global proteome-level changes with corresponding anatomical structures (**Supplemental File S4**), with the link between proteins and specific anatomical structures obtained from the Zebrafish Information Network (ZFIN) database (https://zfin.org/)^86^ based on the traditional in situ hybridization technology. In this work, we focused on the ten major tissue systems in the zebrafish body, including those derived from ectoderm (nervous and integument systems), mesoderm (cardiovascular, musculature, renal, immune, and reproductive systems), and endoderm (respiratory, digestive, and endocrine systems). As shown in **Figure 6 and Supplementary Figure S7**, the expression patterns of tissue-related proteins were similar for different tissue systems, with two groups of proteins offering positive and negative regulations according to their upregulation and downregulation. Interestingly, the emergence of tissue-specific proteins was observed after ZGA during a similar time period, starting from the oblong stage (3.5 hpf) and bursting from the bud stage (10 hpf).

**Figure 6.**
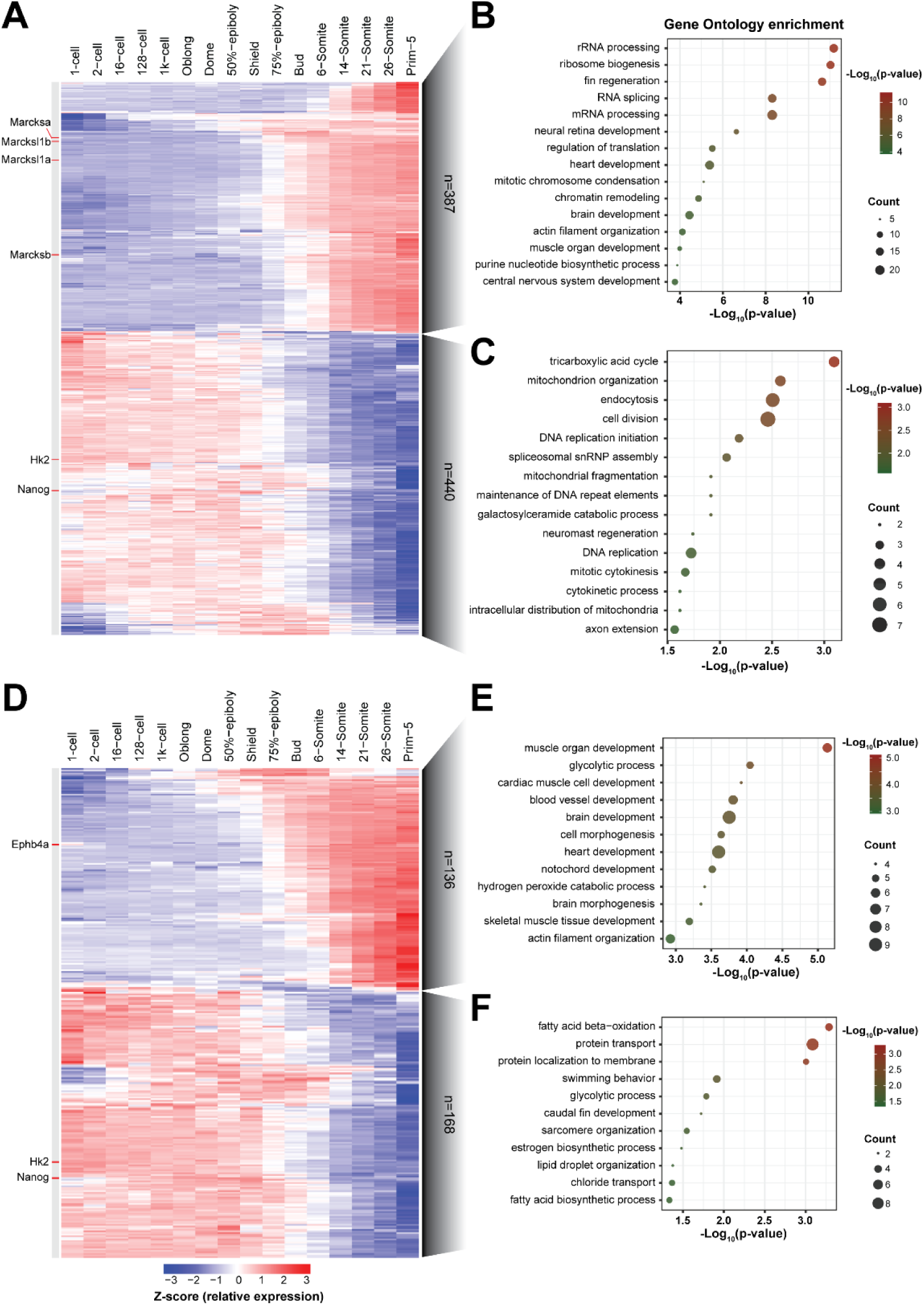
Construction of the tissue resolved developmental trajectories by global proteome expression change data. (A) Heatmap of the expression profiles across the time course for differentially expressed proteins (DEPs) involved in the nervous system. The numbers on the right represent the number of DEPs with an increased or decreased expression trend across embryogenesis. Gene ontology (GO) analysis for DEPs involved in the nervous system, with expression pattern (B) increased and (C) decreased across development, and the enriched biological processes are shown. Adjusted p-values and gene counts are also labeled. (D) Heatmap of the expression profiles across the time course for DEPs involved in the cardiovascular system. The numbers on the right represent the number of DEPs with an increased or decreased expression trend across embryogenesis. GO analysis for DEPs involved in the cardiovascular system, with expression pattern (E) increased and (F) decreased across development, and the enriched biological processes are shown. Adjusted p-values and gene counts are also labeled.

As shown in **Figure 6**, most of the proteins whose expression level declined are involved in fundamental processes such as gene regulation (transcription factor Nanog) and energy metabolism (glycolytic enzyme hexokinase Hk2). Meanwhile, the known tissue-specific proteins, such as the cardiovascular marker protein (receptor protein-tyrosine kinase Ephb4a)^87^, nervous marker protein (myristoylated alanin-rich C-kinase substrate Marcks and Marcks-like 1)^88^, and liver marker protein (fructose-bisphosphate aldolase B Aldob)^89^, which contribute to tissue development, are upregulated across embryogenesis.

Overall, our proteome dataset provides evidence that the TFs and their downstream gene networks are one of the key factors that drive cell fate transition during embryogenesis, while cell types bifurcate into multiple branches at certain time points starting from MZT^90,91^.

### Comparison of mRNA and protein expression patterns during zebrafish embryo development

The transcriptome and proteome typically do not show strong agreement, especially during early embryogenesis^27,92–95^. For example, in our recent study, we observed significant discrepancies between zebrafish proteome and transcriptome profiles during the MZT across four time points^27^. Here, for the first time, we have the opportunity to compare the mRNA and protein expression patterns across a large number of embryonic stages during the first day of the embryo’s life. For this purpose, we integrated our proteomic data with a published transcriptome result^17^, which has served as a time-resolved transcriptional basis for zebrafish genomic changes and has been a helpful resource in systems biology. We used the transcriptome data at 12 embryonic stages close to our proteomic data points from zygote to early pharyngula, including the 1-cell, 2-cell, 128-cell, 1k-cell, dome, 50%-epiboly, shield, 75%-epiboly, 1-4-somite, 14-19-somite, 20-25-somite, and prim-5 stages.

Almost all proteins (2156 out of 2220) differentially expressed in the proteome of zebrafish embryos also had the corresponding mRNA expression data in the transcriptome. Among them, 461 proteins exhibit a strong positive correlation with the RNA expression (p-value<0.05, correlation coefficient>0.6), and 290 proteins show a strong negative correlation (p-value<0.05, correlation coefficient<-0.6, **Supplemental File S6**). Overall, for nearly 80% of the DEPs, the protein and mRNA expression patterns across the 12 time points do not agree, suggesting significant discrepancies between the proteome and transcriptome profiles during the zebrafish early embryogenesis.

The DEPs in each cluster were further grouped into 3 or 4 subclusters based on the expression pattern of the corresponding mRNAs. **Figure 7** shows the data for cluster 1. Protein and mRNA expression dynamics agree well for 32 genes in cluster 1 (subcluster 1-2), **Figure 7A**. These mRNAs are maternal with gradual degradation during MZT, and they are enriched in the L-phenylalanine catabolic process, tetrahydrobiopterin biosynthetic process, P granule, fatty acid elongation, microtubule-based process, fatty acid metabolism, and lysosome. Subclusters 1-1 and 1-3 have 76 and 47 genes, respectively. Proteins and mRNAs show substantial differences in expression dynamics across time points in these subclusters, **Figure 7A**. For example, mRNAs in the subcluster 1-3 have an opposite expression profile to proteins. mRNAs in the subcluster 1-1 have a sudden expression increase during the cleavage period before gradually declining. The discrepancy between mRNAs and proteins is likely due to post-transcriptional and/or post-translational regulations, for example, the control of mRNA degradation, protein translation, and protein degradation. The proteins related to subcluster 1-1 are highly enriched in p-body assembly, stress granule assembly, and oogenesis. The proteins related to the subcluster 1-3 are highly enriched in aerobic respiration, platelet aggregation, mitochondrial electron transport (cytochrome c to oxygen), and oxidative phosphorylation. The results here show that proteins in different subclusters (1-1, 1-2, and 1-3) have unique biological process enrichment profiles, suggesting that the agreement or discrepancy between mRNA and protein expression dynamics may be functionally relevant.

**Figure 7.**
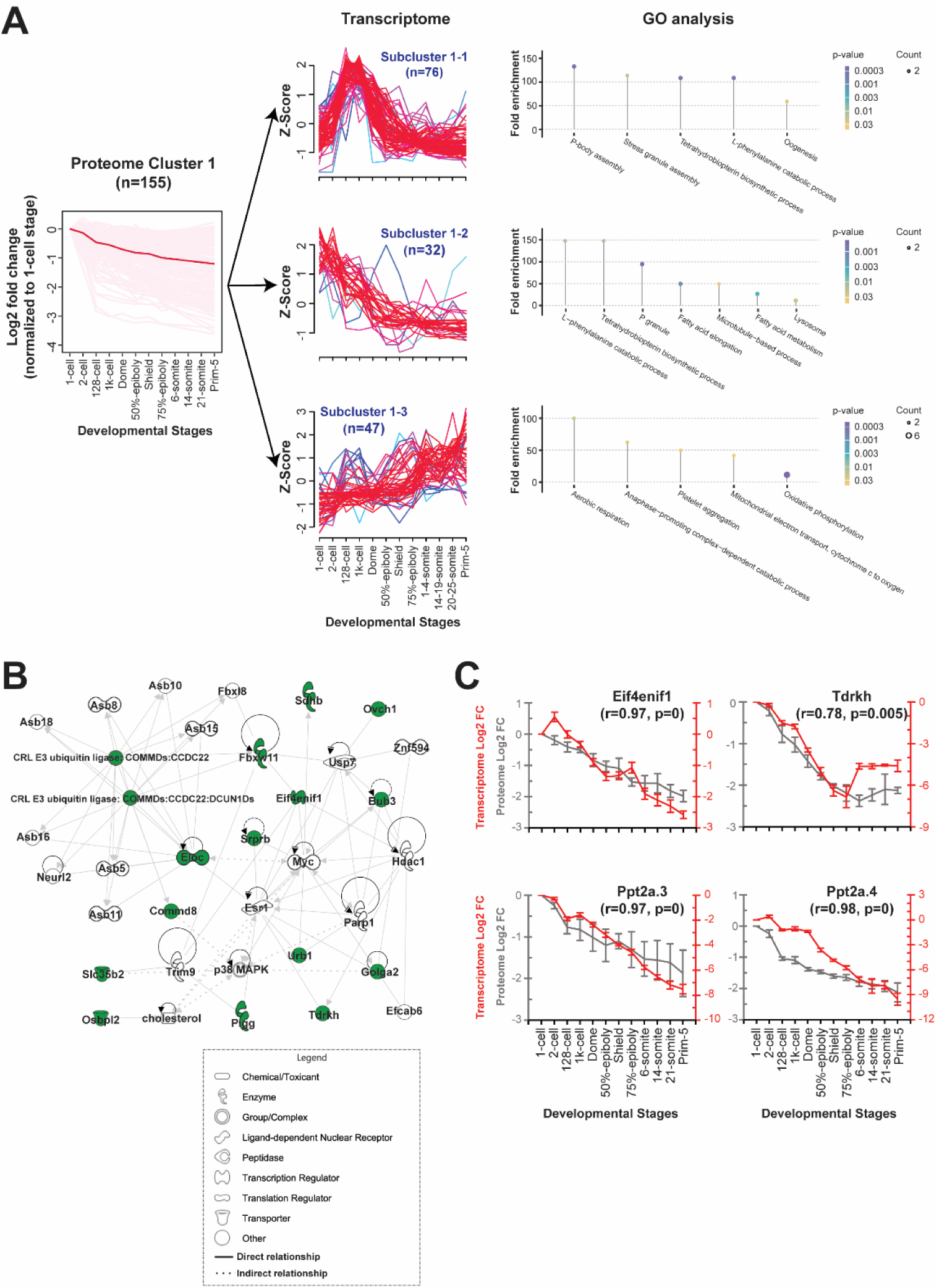
Proteome and transcriptome correlation data for the differentially expressed proteins (DEPs) in cluster 1. (A) The genes of DEPs in cluster 1 are divided into three subclusters based on the dynamics of mRNA abundance across developmental stages. The twelve time points covered by the proteome data are 1-cell, 2-cell, 128-cell, 1k-cell, dome, 50%-epiboly, shield, 75%-epiboly, 6-somite, 14-somite, 21-somite, and prim-5 stages. The time points covered by the transcriptome data are 1-cell, 2-cell, 128-cell, 1k-cell, dome, 50%-epiboly, shield, 75%-epiboly, 1-4-somite, 14-19-somite, 20-25-somite, and prim-5 stages. Gene ontology (GO) analysis for genes in each subcluster was done, and the enriched biological processes are shown. Adjusted p-values and gene counts are also labeled. (B) Ingenuity pathway analysis (IPA) data of proteins in the subcluster 1-2. Proteins marked in green are quantified proteins in this work, and they are involved in cell cycle and protein trafficking pathways. (C) Normalized expression dynamics of mRNAs (red) and proteins (grey) across the 12 time points for four genes in subcluster 1-2. The error bars represent the standard deviations of the fold change from proteome analysis (n=3) and transcriptome analysis (n=5). The correlations between mRNAs and proteins were calculated, with *r* representing the correlation coefficient and p representing the p-value.

We then focused on proteins in the subcluster 1-2, showing agreement between mRNAs and proteins. The QIAGEN Ingenuity Pathway Analysis (IPA) result revealed that the proteins in the subcluster 1-2 were mainly involved in cell cycle and protein transport pathways, **Figure 7B**. **Figure 7C** shows four examples of genes related to the cell cycle from the subcluster 1-2, exhibiting agreement between mRNA and protein expression dynamics across the 12 developmental stages. Specifically, eukaryotic translation initiation factor 4E nuclear import factor 1 (Eif4enif1, correlation coefficient as 0.97), which not only regulates the stability and translation of mRNAs but is also involved in the formation of P-bodies, which store translationally inactive mRNAs^96^, shows equivalent fold changes during zebrafish embryonic development at both mRNA and protein levels. **Figure 8** shows the mRNA and protein correlation data for cluster 4 proteins, which are upregulated from the late gastrula period. Four sub-clusters were generated based on the mRNA data. For subclusters 4-1 and 4-2, nearly 60% of DEPs in cluster 4 are included, and the proteome and transcriptome data agree well, **Figure 8A**. Half of the proteins of subclusters 4-1 and 4-2 are strictly zygotic, and their expression increase is due to the ZGA. Most of the proteins are involved in lipid metabolism, protein synthesis, gene expression, cardiac dilation, and embryonic development pathways, **Figure 8B**, reflecting the main events after ZGA, e.g., organogenesis. More specifically, proteins in subclusters 4-1 and 4-2 are involved in the development of various organs and tissues, e.g., nervous system (Akt1, Cd81a, and Col18a1a), brain (Tubb2b, Tuba1a, Pak2a, and Top2b), heart (Akt1, Actc1a, Myl1, Vclb, Cnn2, and myhz1.2), muscle (HSP90AA1, Myl1, Atp2a1, and myhz1.2), immune system (Rangap1b), eye (Col2a1a and Col18a1a), bone (P3h1 and Cyt1l), skin (Krt17), and tooth (Krt4). The proteins within and between pathways are highly connected, either directly or indirectly, **Figure 8B**, to carry out the complex biological processes involved during the organogenesis period. **Figure 8C** shows four examples of proteins exhibiting consistent expression dynamics between mRNAs and proteins.

**Figure 8.**
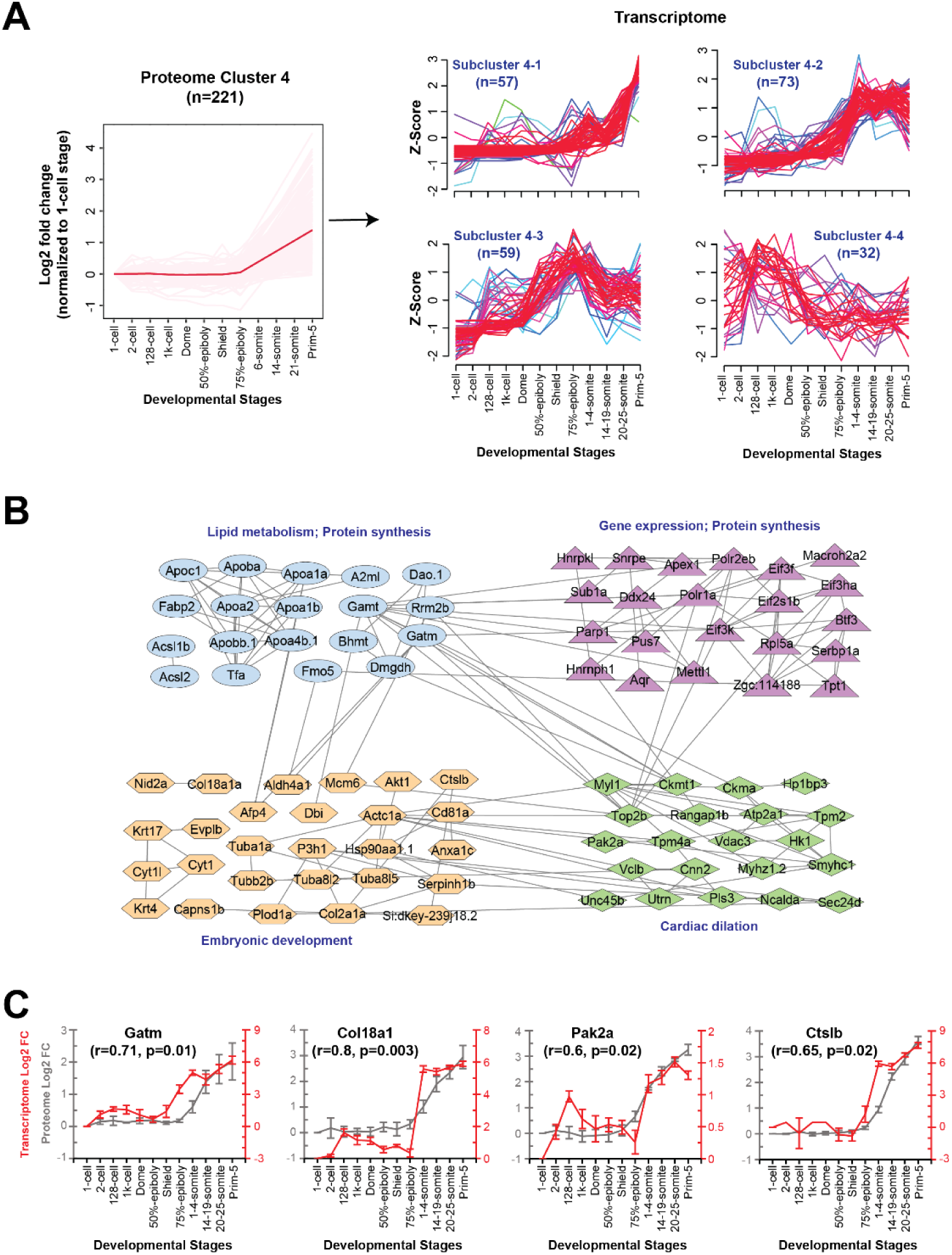
Proteome and transcriptome correlation data for the differentially expressed proteins (DEPs) in cluster 4. (A) The genes of DEPs in cluster 4 are divided into four subclusters based on the dynamics of mRNA abundance across developmental stages. The twelve time points covered by the proteome data and transcriptome data are the same as those in Figure 6. Gene ontology (GO) analysis for genes in each subcluster was done, and the enriched biological processes of genes are shown. Adjusted p-values and gene counts are also labeled. (B) Protein-protein interaction network generated in Cytoscape with DEPs in subclusters 4-1 and 4-2. Different colors represent proteins classified into specific pathways. (C) Normalized expression dynamics of mRNAs (red) and proteins (grey) across the 12 time points for four genes in subclusters 4-1 and 4-2. The error bars represent the standard deviations of the fold change from proteome analysis (n=3) and transcriptome analysis (n=5). The correlations between mRNAs and proteins were calculated, with r representing the correlation coefficient and p representing the p-value.

For subcluster 4-3, the mRNA expression rises before the protein expression increases, followed by a decline in mRNA abundance. The subcluster 4-3 proteins are involved in purine metabolism (pfas, paics, prps1a, impdh1b), eye development (pbx4, mvp, lamc1, lamb1a), brain development (pbx4, mvp, lamc1, lamb1a, aldoaa), and helicase activity (mcm3, dhx9, ddx41, and DEAD-box helicase 41). The protein expression in this subcluster matches the upcoming organogenesis. For proteins in subcluster 4-4, mRNA expression increases before MBT (1k-cell stage), followed by a substantial drop in early gastrula and a stable expression through gastrula and segmentation periods. The proteins are involved in kinase activity (rock1, thap12a, camk2g1, cmpk, dtymk, and insra). The discrepancy between mRNAs and proteins in those two subclusters further indicates the diversity of post-transcriptional and post-translational regulations during early embryogenesis.

**Figures S8-S10** show the mRNA and protein correlation data for protein clusters 2 and 3, 5 and 6, and 7 and 8, respectively. Overall, for the majority of the proteins in those clusters, the mRNA and protein expression profiles do not agree, and the enriched biological processes in the subclusters are drastically different, indicating that the expression discrepancy between mRNAs and proteins should be functionally driven.

We then focused on protein cluster 5, containing the highest number of DEPs, **Figure S9A**. For subclusters 5-1 and 5-2, the mRNA and protein expression agree well for the cleavage, blastula, and gastrula periods, with a gradual increase. However, mRNA abundance declines (subcluster 5-1) or remains relatively constant (subcluster 5-2) throughout the segmentation period, whereas protein abundance increases rapidly. The GO enrichment data show that proteins in subclusters 5-1 and 5-2 are highly relevant to organogenesis, e.g., neural crest formation, cardiac muscle cell development, hematopoietic stem cell differentiation, heart development, brain development, neural retina development, blood vessel development, and fin regeneration. Subcluster 5-3 proteins are enriched in mRNA processing, **Figures S9A** and **S11A**, and their mRNAs exhibit opposite expression profiles to the proteins, with a gradual decrease in abundance throughout early development, likely due to maternal mRNA degradation. Similar to subcluster 5-3, subcluster 6-1 proteins also exhibit opposite expression profiles at the mRNA level. mRNAs of subcluster 6-1 have an abundance increase throughout the studied developmental period due to likely ZGA, whereas proteins have a gradual decline in expression. Proteins in subcluster 6-1 are involved in proteolysis, protein transport, protein N-linked glycosylation, and oxidative phosphorylation, **Figures S9B** and **S11B**.

## Conclusion

The ability to define how protein abundance changes during organogenesis is central to our understanding of animal development and human disease. Collectively, this work provides a comprehensive analysis of the dynamic protein expression across 16 stages throughout zebrafish embryogenesis for insight into these developmental processes. The large number of stages allows us to cluster 4418 proteins by their expression profile across all of development, discovering a plethora of key proteins regulating zebrafish embryo development. The zinc finger transcription factors in chromosome 4 are highly expressed during ZGA. As two distinct layers of gene regulation, we found a strong discordance between the proteome and transcriptome datasets. Interestingly, we also investigated the strong correlation for metabolism, cytoskeletal organization, and translation machinery proteins between protein and mRNA levels. This atlas provides a valuable resource for unraveling uncharacterized layers of gene regulation during zebrafish organogenesis.

## Acknowledgments

## Funding

The work was funded by the National Institute of General Medical Sciences (NIGMS) through grant R01GM125991. We also thank the support from the National Cancer Institute (NCI) through the grant R01CA247863, and the National Institute of General Medical Sciences (NIGMS) through grant R35GM153479. Sun also thanks the support from the National Science Foundation (CAREER Award, Grant DBI1846913). We thank MSU AgBioResearch and Michigan State University for access to the QIAGEN Ingenuity Pathway Analysis (IPA) platform.

## Author contributions

FF performed the quantitative proteomics experiments. FF performed all the data analysis and made the first draft of the manuscript. YY performed the statistical analysis. ARB and XL built a website for distributing quantitative proteomics datasets. BP and JBC maintained the zebrafish animals and helped with the embryo collections. LS conceived the original idea and supervised the project. All authors provided comments and contributed to the final manuscript.

## Competing interests

The Authors declare that they have no competing interests.

## Data and materials availability

All data needed to evaluate the conclusions in the paper are present in the paper and/or the Supplementary Materials. The MS raw data have been deposited to the ProteomeXchange Consortium via the PRIDE^97^ partner repository with the dataset identifier PXD070423. We have built a website (https://www.toppic.org/software/zebrafishdb/zf_protein.html) to facilitate developmental biologists to get access to the quantitative proteomics dataset.

## Notes

### Competing Interest Statement

The authors have declared no competing interest.

## Reference

1. Tadros, W., and Lipshitz, H.D. (2009). The maternal-to-zygotic transition: a play in two acts. Development 136, 3033–3042. 10.1242/dev.033183.

2. Vastenhouw, N.L., Cao, W.X., and Lipshitz, H.D. (2019). The maternal-to-zygotic transition revisited. Development 146, dev161471. 10.1242/dev.161471.

3. Shahbazi, M.N. (2020). Mechanisms of human embryo development: from cell fate to tissue shape and back. Development 147, dev190629. 10.1242/dev.190629.

4. Veldman, M.B., and Lin, S. (2008). Zebrafish as a Developmental Model Organism for Pediatric Research. Pediatr Res 64, 470–476. 10.1203/PDR.0b013e318186e609.

5. White, R.M., Sessa, A., Burke, C., Bowman, T., LeBlanc, J., Ceol, C., Bourque, C., Dovey, M., Goessling, W., Burns, C.E., et al. (2008). Transparent adult zebrafish as a tool for in vivo transplantation analysis. Cell Stem Cell 2, 183–189. 10.1016/j.stem.2007.11.002.

6. Kimmel, C.B., Ballard, W.W., Kimmel, S.R., Ullmann, B., and Schilling, T.F. (1995). Stages of embryonic development of the zebrafish. Dev Dyn 203, 253–310. 10.1002/aja.1002030302.

7. Avdesh, A., Chen, M., Martin-Iverson, M.T., Mondal, A., Ong, D., Rainey-Smith, S., Taddei, K., Lardelli, M., Groth, D.M., Verdile, G., et al. (2012). Regular Care and Maintenance of a Zebrafish (Danio rerio) Laboratory: An Introduction. J Vis Exp, 4196. 10.3791/4196.

8. Howe, K., Clark, M.D., Torroja, C.F., Torrance, J., Berthelot, C., Muffato, M., Collins, J.E., Humphray, S., McLaren, K., Matthews, L., et al. (2013). The zebrafish reference genome sequence and its relationship to the human genome. Nature 496, 498–503. 10.1038/nature12111.

9. Yashaswini, C., Kiran, N.S., and Chatterjee, A. (2025). Zebrafish navigating the metabolic maze: insights into human disease - assets, challenges and future implications. J Diabetes Metab Disord 24, 3. 10.1007/s40200-024-01539-8.

10. Asharani, P.V., Keupp, K., Semler, O., Wang, W., Li, Y., Thiele, H., Yigit, G., Pohl, E., Becker, J., Frommolt, P., et al. (2012). Attenuated BMP1 function compromises osteogenesis, leading to bone fragility in humans and zebrafish. Am J Hum Genet 90, 661–674. 10.1016/j.ajhg.2012.02.026.

11. Fiedler, I.A.K., Schmidt, F.N., Wölfel, E.M., Plumeyer, C., Milovanovic, P., Gioia, R., Tonelli, F., Bale, H.A., Jähn, K., Besio, R., et al. (2018). Severely Impaired Bone Material Quality in Chihuahua Zebrafish Resembles Classical Dominant Human Osteogenesis Imperfecta. J Bone Miner Res 33, 1489–1499. 10.1002/jbmr.3445.

12. LaBonty, M., Pray, N., and Yelick, P.C. (2017). A Zebrafish Model of Human Fibrodysplasia Ossificans Progressiva. Zebrafish 14, 293–304. 10.1089/zeb.2016.1398.

13. Busse, B., Galloway, J.L., Gray, R.S., Harris, M.P., and Kwon, R.Y. (2020). Zebrafish: An Emerging Model for Orthopedic Research. J Orthop Res 38, 925–936. 10.1002/jor.24539.

14. Lee, M.T., Bonneau, A.R., Takacs, C.M., Bazzini, A.A., DiVito, K.R., Fleming, E.S., and Giraldez, A.J. (2013). Nanog, Pou5f1 and SoxB1 activate zygotic gene expression during the maternal-to-zygotic transition. Nature 503, 360–364. 10.1038/nature12632.

15. Leichsenring, M., Maes, J., Mössner, R., Driever, W., and Onichtchouk, D. (2013). Pou5f1 transcription factor controls zygotic gene activation in vertebrates. Science 341, 1005–1009. 10.1126/science.1242527.

16. Pálfy, M., Schulze, G., Valen, E., and Vastenhouw, N.L. (2020). Chromatin accessibility established by Pou5f3, Sox19b and Nanog primes genes for activity during zebrafish genome activation. PLoS Genet 16, e1008546. 10.1371/journal.pgen.1008546.

17. White, R.J., Collins, J.E., Sealy, I.M., Wali, N., Dooley, C.M., Digby, Z., Stemple, D.L., Murphy, D.N., Billis, K., Hourlier, T., et al. (2017). A high-resolution mRNA expression time course of embryonic development in zebrafish. Elife 6, e30860. 10.7554/eLife.30860.

18. Reimão-Pinto, M.M., Behrens, A., Forcelloni, S., Fröhlich, K., Kaya, S., and Nedialkova, D.D. (2024). The dynamics and functional impact of tRNA repertoires during early embryogenesis in zebrafish. EMBO J 43, 5747–5779. 10.1038/s44318-024-00265-4.

19. Tay, T.L., Lin, Q., Seow, T.K., Tan, K.H., Hew, C.L., and Gong, Z. (2006). Proteomic analysis of protein profiles during early development of the zebrafish, Danio rerio. Proteomics 6, 3176–3188. 10.1002/pmic.200600030.

20. Lucitt, M.B., Price, T.S., Pizarro, A., Wu, W., Yocum, A.K., Seiler, C., Pack, M.A., Blair, I.A., Fitzgerald, G.A., and Grosser, T. (2008). Analysis of the zebrafish proteome during embryonic development. Mol Cell Proteomics 7, 981–994. 10.1074/mcp.M700382-MCP200.

21. Lößner, C., Wee, S., Ler, S.G., Li, R.H.X., Carney, T., Blackstock, W., and Gunaratne, J. (2012). Expanding the zebrafish embryo proteome using multiple fractionation approaches and tandem mass spectrometry. Proteomics 12, 1879–1882. 10.1002/pmic.201100576.

22. Alli Shaik, A., Wee, S., Li, R.H.X., Li, Z., Carney, T.J., Mathavan, S., and Gunaratne, J. (2014). Functional mapping of the zebrafish early embryo proteome and transcriptome. J Proteome Res 13, 5536–5550. 10.1021/pr5005136.

23. Lin, Y., Chen, Y., Yang, X., Xu, D., and Liang, S. (2009). Proteome analysis of a single zebrafish embryo using three different digestion strategies coupled with liquid chromatography-tandem mass spectrometry. Anal Biochem 394, 177–185. 10.1016/j.ab.2009.07.034.

24. Purushothaman, K., Das, P.P., Presslauer, C., Lim, T.K., Johansen, S.D., Lin, Q., and Babiak, I. (2019). Proteomics Analysis of Early Developmental Stages of Zebrafish Embryos. Int J Mol Sci 20, 6359. 10.3390/ijms20246359.

25. Kwon, O.K., Kim, S.J., Lee, Y.-M., Lee, Y.-H., Bae, Y.-S., Kim, J.Y., Peng, X., Cheng, Z., Zhao, Y., and Lee, S. (2016). Global analysis of phosphoproteome dynamics in embryonic development of zebrafish (Danio rerio). Proteomics 16, 136–149. 10.1002/pmic.201500017.

26. da Silva Pescador, G., Baia Amaral, D., Varberg, J.M., Zhang, Y., Hao, Y., Florens, L., and Bazzini, A.A. (2024). Protein profiling of zebrafish embryos unmasks regulatory layers during early embryogenesis. Cell Rep 43, 114769. 10.1016/j.celrep.2024.114769.

27. Fang, F., Chen, D., Basharat, A.R., Poulos, W., Wang, Q., Cibelli, J.B., Liu, X., and Sun, L. (2024). Quantitative proteomics reveals the dynamic proteome landscape of zebrafish embryos during the maternal-to-zygotic transition. iScience 27, 109944. 10.1016/j.isci.2024.109944.

28. Yan, J., Ding, Y., Peng, Z., Qin, L., Gu, J., and Wan, C. (2023). Systematic Proteomics Study on the Embryonic Development of Danio rerio. J. Proteome Res. 22, 2814–2826. 10.1021/acs.jproteome.3c00056.

29. Cox, J., and Mann, M. (2008). MaxQuant enables high peptide identification rates, individualized p.p.b.-range mass accuracies and proteome-wide protein quantification. Nat Biotechnol 26, 1367–1372. 10.1038/nbt.1511.

30. Tyanova, S., Temu, T., Sinitcyn, P., Carlson, A., Hein, M.Y., Geiger, T., Mann, M., and Cox, J. (2016). The Perseus computational platform for comprehensive analysis of (prote)omics data. Nat Methods 13, 731–740. 10.1038/nmeth.3901.

31. Chawade, A., Alexandersson, E., and Levander, F. (2014). Normalyzer: a tool for rapid evaluation of normalization methods for omics data sets. J Proteome Res 13, 3114–3120. 10.1021/pr401264n.

32. Conesa, A., Nueda, M.J., Ferrer, A., and Talón, M. (2006). maSigPro: a method to identify significantly differential expression profiles in time-course microarray experiments. Bioinformatics 22, 1096–1102. 10.1093/bioinformatics/btl056.

33. Zhang, J. (2025). ClusterGVis: One-Step to Cluster and Visualize Gene Expression Data. 10.32614/CRAN.package.ClusterGVis https://doi.org/10.32614/CRAN.package.ClusterGVis.

34. Venables, W.N., and Ripley, B.D. (2002). Modern Applied Statistics with S (Springer) 10.1007/978-0-387-21706-2.

35. Wickham, H. (2009). ggplot2: Elegant Graphics for Data Analysis (Springer) 10.1007/978-0-387-98141-3.

36. Gu, Z., Gu, L., Eils, R., Schlesner, M., and Brors, B. (2014). circlize Implements and enhances circular visualization in R. Bioinformatics 30, 2811–2812. 10.1093/bioinformatics/btu393.

37. Sherman, B.T., Hao, M., Qiu, J., Jiao, X., Baseler, M.W., Lane, H.C., Imamichi, T., and Chang, W. (2022). DAVID: a web server for functional enrichment analysis and functional annotation of gene lists (2021 update). Nucleic Acids Res 50, W216–W221. 10.1093/nar/gkac194.

38. Huang, D.W., Sherman, B.T., and Lempicki, R.A. (2009). Systematic and integrative analysis of large gene lists using DAVID bioinformatics resources. Nat Protoc 4, 44–57. 10.1038/nprot.2008.211.

39. Supek, F., Bošnjak, M., Škunca, N., and Šmuc, T. (2011). REVIGO summarizes and visualizes long lists of gene ontology terms. PLoS One 6, e21800. 10.1371/journal.pone.0021800.

40. Szklarczyk, D., Kirsch, R., Koutrouli, M., Nastou, K., Mehryary, F., Hachilif, R., Gable, A.L., Fang, T., Doncheva, N.T., Pyysalo, S., et al. (2023). The STRING database in 2023: protein-protein association networks and functional enrichment analyses for any sequenced genome of interest. Nucleic Acids Res 51, D638–D646. 10.1093/nar/gkac1000.

41. Demchak, B., Hull, T., Reich, M., Liefeld, T., Smoot, M., Ideker, T., and Mesirov, J.P. (2014). Cytoscape: the network visualization tool for GenomeSpace workflows. F1000Res 3, 151. 10.12688/f1000research.4492.2.

42. Best, D.J., and Roberts, D.E. (1975). Algorithm AS 89: The Upper Tail Probabilities of Spearman’s Rho. Journal of the Royal Statistical Society. Series C (Applied Statistics) 24, 377–379. 10.2307/2347111.

43. Link, V., Shevchenko, A., and Heisenberg, C.-P. (2006). Proteomics of early zebrafish embryos. BMC Dev Biol 6, 1. 10.1186/1471-213X-6-1.

44. Rahlouni, F., Szarka, S., Shulaev, V., and Prokai, L. (2015). A Survey of the Impact of Deyolking on Biological Processes Covered by Shotgun Proteomic Analyses of Zebrafish Embryos. Zebrafish 12, 398–407. 10.1089/zeb.2015.1121.

45. Thompson, A., Wölmer, N., Koncarevic, S., Selzer, S., Böhm, G., Legner, H., Schmid, P., Kienle, S., Penning, P., Höhle, C., et al. (2019). TMTpro: Design, Synthesis, and Initial Evaluation of a Proline-Based Isobaric 16-Plex Tandem Mass Tag Reagent Set. Anal Chem 91, 15941–15950. 10.1021/acs.analchem.9b04474.

46. Swearingen, K.E., and Moritz, R.L. (2012). High-field asymmetric waveform ion mobility spectrometry for mass spectrometry-based proteomics. Expert Rev Proteomics 9, 505–517. 10.1586/epr.12.50.

47. Schnabel, D., Castillo-Robles, J., and Lomeli, H. (2019). Protein Purification and Western Blot Detection from Single Zebrafish Embryo. Zebrafish 16, 505–507. 10.1089/zeb.2019.1761.

48. He, M., Zhang, R., Jiao, S., Zhang, F., Ye, D., Wang, H., and Sun, Y. (2020). Nanog safeguards early embryogenesis against global activation of maternal β-catenin activity by interfering with TCF factors. PLoS Biol 18, e3000561. 10.1371/journal.pbio.3000561.

49. Castillo-Robles, J., Ramírez, L., Spaink, H.P., and Lomelí, H. (2018). smarce1 mutants have a defective endocardium and an increased expression of cardiac transcription factors in zebrafish. Sci Rep 8, 15369. 10.1038/s41598-018-33746-8.

50. Krens, S.G., Corredor-Adámez, M., He, S., Snaar-Jagalska, B.E., and Spaink, H.P. (2008). ERK1 and ERK2 MAPK are key regulators of distinct gene sets in zebrafish embryogenesis. BMC Genomics 9, 196. 10.1186/1471-2164-9-196.

51. Theocharidis, A., van Dongen, S., Enright, A.J., and Freeman, T.C. (2009). Network visualization and analysis of gene expression data using BioLayout Express(3D). Nat Protoc 4, 1535–1550. 10.1038/nprot.2009.177.

52. Bhat, P., Cabrera-Quio, L.E., Herzog, V.A., Fasching, N., Pauli, A., and Ameres, S.L. (2023). SLAMseq resolves the kinetics of maternal and zygotic gene expression during early zebrafish embryogenesis. Cell Rep 42, 112070. 10.1016/j.celrep.2023.112070.

53. Lu, X., Li, J.M., Elemento, O., Tavazoie, S., and Wieschaus, E.F. (2009). Coupling of zygotic transcription to mitotic control at the Drosophila mid-blastula transition. Development 136, 2101–2110. 10.1242/dev.034421.

54. Adar-Levor, S., Nachmias, D., Gal-Oz, S.T., Jahn, Y.M., Peyrieras, N., Zaritsky, A., Birnbaum, R.Y., and Elia, N. (2021). Cytokinetic abscission is part of the midblastula transition in early zebrafish embryogenesis. Proc Natl Acad Sci U S A 118, e2021210118. 10.1073/pnas.2021210118.

55. Sophos, N.A., and Vasiliou, V. (2003). Aldehyde dehydrogenase gene superfamily: the 2002 update. Chem Biol Interact 143*–144*, 5–22. 10.1016/s0009-2797(02)00163-1.

56. Brunsdon, H., Brombin, A., Peterson, S., Postlethwait, J.H., and Patton, E.E. (2022). Aldh2 is a lineage-specific metabolic gatekeeper in melanocyte stem cells. Development 149, dev200277. 10.1242/dev.200277.

57. Furukawa, F., Aoyagi, A., Sano, K., Sameshima, K., Goto, M., Tseng, Y.-C., Ikeda, D., Lin, C.-C., Uchida, K., Okumura, S.-I., et al. (2024). Gluconeogenesis in the extraembryonic yolk syncytial layer of the zebrafish embryo. PNAS Nexus 3, pgae125. 10.1093/pnasnexus/pgae125.

58. Duque Lasio, M.L., Lehman, A.N., Ahmad, A., and Bedoyan, J.K. (1993). Pyruvate Carboxylase Deficiency. In GeneReviews®, M. P. Adam, S. Bick, G. M. Mirzaa, R. A. Pagon, S. E. Wallace, and A. Amemiya, eds. (University of Washington, Seattle).

59. Fischer, S., Prykhozhij, S., Rau, M.J., and Neumann, C.J. (2007). Mutation of zebrafish caf-1b results in S phase arrest, defective differentiation, and p53-mediated apoptosis during organogenesis. Cell Cycle 6, 2962–2969. 10.4161/cc.6.23.4950.

60. Chen, Z., Tan, J.L.H., Ingouff, M., Sundaresan, V., and Berger, F. (2008). Chromatin assembly factor 1 regulates the cell cycle but not cell fate during male gametogenesis in Arabidopsis thaliana. Development 135, 65–73. 10.1242/dev.010108.

61. Zeng, C., Han, S., Pan, Y., Huang, Z., Zhang, B., and Zhang, B. (2024). Revisiting the chaperonin T-complex protein-1 ring complex in human health and disease: A proteostasis modulator and beyond. Clin Transl Med 14, e1592. 10.1002/ctm2.1592.

62. Zamir, E., Kam, Z., and Yarden, A. (1997). Transcription-dependent induction of G1 phase during the zebra fish midblastula transition. Mol Cell Biol 17, 529–536. 10.1128/MCB.17.2.529.

63. Collart, C., Owens, N.D.L., Bhaw-Rosun, L., Cooper, B., De Domenico, E., Patrushev, I., Sesay, A.K., Smith, J.N., Smith, J.C., and Gilchrist, M.J. (2014). High-resolution analysis of gene activity during the Xenopus mid-blastula transition. Development 141, 1927–1939. 10.1242/dev.102012.

64. Chemnitz, J., Pieper, D., Stich, L., Schumacher, U., Balabanov, S., Spohn, M., Grundhoff, A., Steinkasserer, A., Hauber, J., and Zinser, E. (2019). The acidic protein rich in leucines Anp32b is an immunomodulator of inflammation in mice. Sci Rep 9, 4853. 10.1038/s41598-019-41269-z.

65. Christov, C.P., Dingwell, K.S., Skehel, M., Wilkes, H.S., Sale, J.E., Smith, J.C., and Krude, T. (2018). A NuRD Complex from Xenopus laevis Eggs Is Essential for DNA Replication during Early Embryogenesis. Cell Rep 22, 2265–2278. 10.1016/j.celrep.2018.02.015.

66. Montarras, D., Fiszman, M.Y., and Gros, F. (1982). Changes in tropomyosin during development of chick embryonic skeletal muscles in vivo and during differentiation of chick muscle cells in vitro. J Biol Chem 257, 545–548.

67. Kulmanov, M., Guzmán-Vega, F.J., Duek Roggli, P., Lane, L., Arold, S.T., and Hoehndorf, R. (2024). Protein function prediction as approximate semantic entailment. Nat Mach Intell 6, 220–228. 10.1038/s42256-024-00795-w.

68. Ringuette, M.J., Chamberlin, M.E., Baur, A.W., Sobieski, D.A., and Dean, J. (1988). Molecular analysis of cDNA coding for ZP3, a sperm binding protein of the mouse zona pellucida. Dev Biol 127, 287–295. 10.1016/0012-1606(88)90315-6.

69. Zhang, S., and Cui, P. (2014). Complement system in zebrafish. Dev Comp Immunol 46, 3–10. 10.1016/j.dci.2014.01.010.

70. Nassari, S., Del Olmo, T., and Jean, S. (2020). Rabs in Signaling and Embryonic Development. Int J Mol Sci 21, 1064. 10.3390/ijms21031064.

71. Kenyon, E.J., Campos, I., Bull, J.C., Williams, P.H., Stemple, D.L., and Clark, M.D. (2015). Zebrafish Rab5 proteins and a role for Rab5ab in nodal signalling. Dev Biol 397, 212–224. 10.1016/j.ydbio.2014.11.007.

72. Zhang, H., Gao, Y., Qian, P., Dong, Z., Hao, W., Liu, D., and Duan, X. (2019). Expression analysis of Rab11 during zebrafish embryonic development. BMC Dev Biol 19, 25. 10.1186/s12861-019-0207-7.

73. Li, X., Zhou, W., Li, X., Gao, M., Ji, S., Tian, W., Ji, G., Du, J., and Hao, A. (2019). SOX19b regulates the premature neuronal differentiation of neural stem cells through EZH2-mediated histone methylation in neural tube development of zebrafish. Stem Cell Res Ther 10, 389. 10.1186/s13287-019-1495-3.

74. Ding, D., Bergmaier, P., Sachs, P., Klangwart, M., Rückert, T., Bartels, N., Demmers, J., Dekker, M., Poot, R.A., and Mermoud, J.E. (2018). The CUE1 domain of the SNF2-like chromatin remodeler SMARCAD1 mediates its association with KRAB-associated protein 1 (KAP1) and KAP1 target genes. J Biol Chem 293, 2711–2724. 10.1074/jbc.RA117.000959.

75. Xiao, S., Lu, J., Sridhar, B., Cao, X., Yu, P., Zhao, T., Chen, C.-C., McDee, D., Sloofman, L., Wang, Y., et al. (2017). SMARCAD1 Contributes to the Regulation of Naive Pluripotency by Interacting with Histone Citrullination. Cell Rep 18, 3117–3128. 10.1016/j.celrep.2017.02.070.

76. Efroni, S., Duttagupta, R., Cheng, J., Dehghani, H., Hoeppner, D.J., Dash, C., Bazett-Jones, D.P., Le Grice, S., McKay, R.D.G., Buetow, K.H., et al. (2008). Global transcription in pluripotent embryonic stem cells. Cell Stem Cell 2, 437–447. 10.1016/j.stem.2008.03.021.

77. Moleri, S., Cappellano, G., Gaudenzi, G., Cermenati, S., Cotelli, F., Horner, D.S., and Beltrame, M. (2011). The HMGB protein gene family in zebrafish: Evolution and embryonic expression patterns. Gene Expr Patterns 11, 3–11. 10.1016/j.gep.2010.08.006.

78. Zhao, Y., Yang, Z., Wu, J., Wu, R., Keshipeddy, S.K., Wright, D., and Wang, L. (2017). High-mobility-group protein 2 regulated by microRNA-127 and small heterodimer partner modulates pluripotency of mouse embryonic stem cells and liver tumor initiating cells. Hepatol Commun 1, 816–830. 10.1002/hep4.1086.

79. Rivera, C., Lee, H.-G., Lappala, A., Wang, D., Noches, V., Olivares-Costa, M., Sjöberg-Herrera, M., Lee, J.T., and Andrés, M.E. (2022). Unveiling RCOR1 as a rheostat at transcriptionally permissive chromatin. Nat Commun 13, 1550. 10.1038/s41467-022-29261-0.

80. Tai, H.H., Geisterfer, M., Bell, J.C., Moniwa, M., Davie, J.R., Boucher, L., and McBurney, M.W. (2003). CHD1 associates with NCoR and histone deacetylase as well as with RNA splicing proteins. Biochem Biophys Res Commun 308, 170–176. 10.1016/s0006-291x(03)01354-8.

81. Hennig, B.P., Bendrin, K., Zhou, Y., and Fischer, T. (2012). Chd1 chromatin remodelers maintain nucleosome organization and repress cryptic transcription. EMBO Rep 13, 997–1003. 10.1038/embor.2012.146.

82. Kok, F.O., Taibi, A., Wanner, S.J., Xie, X., Moravec, C.E., Love, C.E., Prince, V.E., Mumm, J.S., and Sirotkin, H.I. (2012). Zebrafish rest regulates developmental gene expression but not neurogenesis. Development 139, 3838–3848. 10.1242/dev.080994.

83. Nolte, H., Konzer, A., Ruhs, A., Jungblut, B., Braun, T., and Krüger, M. (2014). Global protein expression profiling of zebrafish organs based on in vivo incorporation of stable isotopes. J Proteome Res 13, 2162–2174. 10.1021/pr5000335.

84. Kelkar, D.S., Provost, E., Chaerkady, R., Muthusamy, B., Manda, S.S., Subbannayya, T., Selvan, L.D.N., Wang, C.-H., Datta, K.K., Woo, S., et al. (2014). Annotation of the zebrafish genome through an integrated transcriptomic and proteomic analysis. Mol Cell Proteomics 13, 3184–3198. 10.1074/mcp.M114.038299.

85. Abramsson, A., Westman-Brinkmalm, A., Pannee, J., Gustavsson, M., von Otter, M., Blennow, K., Brinkmalm, G., Kettunen, P., and Zetterberg, H. (2010). Proteomics profiling of single organs from individual adult zebrafish. Zebrafish 7, 161–168. 10.1089/zeb.2009.0644.

86. Bradford, Y., Conlin, T., Dunn, N., Fashena, D., Frazer, K., Howe, D.G., Knight, J., Mani, P., Martin, R., Moxon, S.A.T., et al. (2011). ZFIN: enhancements and updates to the Zebrafish Model Organism Database. Nucleic Acids Res 39, D822–829. 10.1093/nar/gkq1077.

87. Kawasaki, J., Aegerter, S., Fevurly, R.D., Mammoto, A., Mammoto, T., Sahin, M., Mably, J.D., Fishman, S.J., and Chan, J. (2014). RASA1 functions in EPHB4 signaling pathway to suppress endothelial mTORC1 activity. J Clin Invest 124, 2774–2784. 10.1172/JCI67084.

88. Prieto, D., and Zolessi, F.R. (2017). Functional Diversification of the Four MARCKS Family Members in Zebrafish Neural Development. J Exp Zool B Mol Dev Evol 328, 119–138. 10.1002/jez.b.22691.

89. Shiokawa, K., Kajita, E., Hara, H., Yatsuki, H., and Hori, K. (2002). A developmental biological study of aldolase gene expression in Xenopus laevis. Cell Res 12, 85–96. 10.1038/sj.cr.7290114.

90. Wilkinson, A.C., Nakauchi, H., and Göttgens, B. (2017). Mammalian Transcription Factor Networks: Recent Advances in Interrogating Biological Complexity. Cell Syst 5, 319–331. 10.1016/j.cels.2017.07.004.

91. Liu, C., Li, R., Li, Y., Lin, X., Zhao, K., Liu, Q., Wang, S., Yang, X., Shi, X., Ma, Y., et al. (2022). Spatiotemporal mapping of gene expression landscapes and developmental trajectories during zebrafish embryogenesis. Dev Cell 57, 1284–1298.e5. 10.1016/j.devcel.2022.04.009.

92. Peshkin, L., Wühr, M., Pearl, E., Haas, W., Freeman, R.M., Gerhart, J.C., Klein, A.M., Horb, M., Gygi, S.P., and Kirschner, M.W. (2015). On the Relationship of Protein and mRNA Dynamics in Vertebrate Embryonic Development. Dev Cell 35, 383–394. 10.1016/j.devcel.2015.10.010.

93. Griffin, T.J., Gygi, S.P., Ideker, T., Rist, B., Eng, J., Hood, L., and Aebersold, R. (2002). Complementary profiling of gene expression at the transcriptome and proteome levels in Saccharomyces cerevisiae. Mol Cell Proteomics 1, 323–333. 10.1074/mcp.m200001-mcp200.

94. Tian, Q., Stepaniants, S.B., Mao, M., Weng, L., Feetham, M.C., Doyle, M.J., Yi, E.C., Dai, H., Thorsson, V., Eng, J., et al. (2004). Integrated genomic and proteomic analyses of gene expression in Mammalian cells. Mol Cell Proteomics 3, 960–969. 10.1074/mcp.M400055-MCP200.

95. Liu, Y., Beyer, A., and Aebersold, R. (2016). On the Dependency of Cellular Protein Levels on mRNA Abundance. Cell 165, 535–550. 10.1016/j.cell.2016.03.014.

96. Ding, Y., He, Z., Sha, Y., Kee, K., and Li, L. (2023). Eif4enif1 haploinsufficiency disrupts oocyte mitochondrial dynamics and leads to subfertility. Development 150, dev202151. 10.1242/dev.202151.

97. Deutsch, E.W., Bandeira, N., Perez-Riverol, Y., Sharma, V., Carver, J.J., Mendoza, L., Kundu, D.J., Wang, S., Bandla, C., Kamatchinathan, S., et al. (2023). The ProteomeXchange consortium at 10 years: 2023 update. Nucleic Acids Res 51, D1539–D1548. 10.1093/nar/gkac1040.

