## Supplementary File S1 for "Uncovering zebrafish embryonic proteome dynamics across 16 time points during the first 24 hours of development"

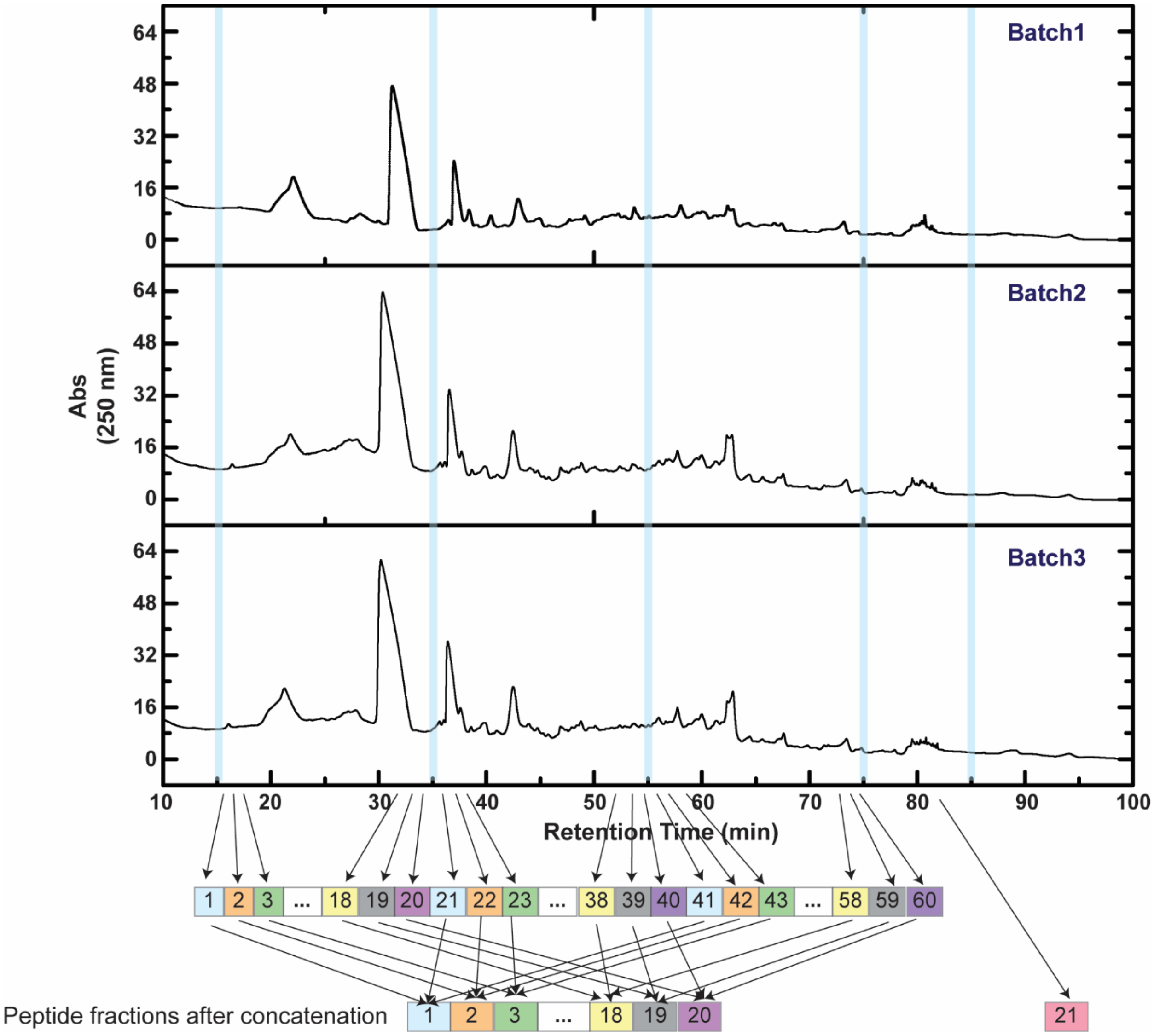


**Figure S1**. Reproducibility of the separation of TMT-labeled zebrafish embryo tryptic peptides with first-dimension high-pH reversed-phase liquid chromatography (RPLC) in biological triplicates. The bottom is a schematic depicting the concatenation strategy applied to the high-pH RPLC separation.

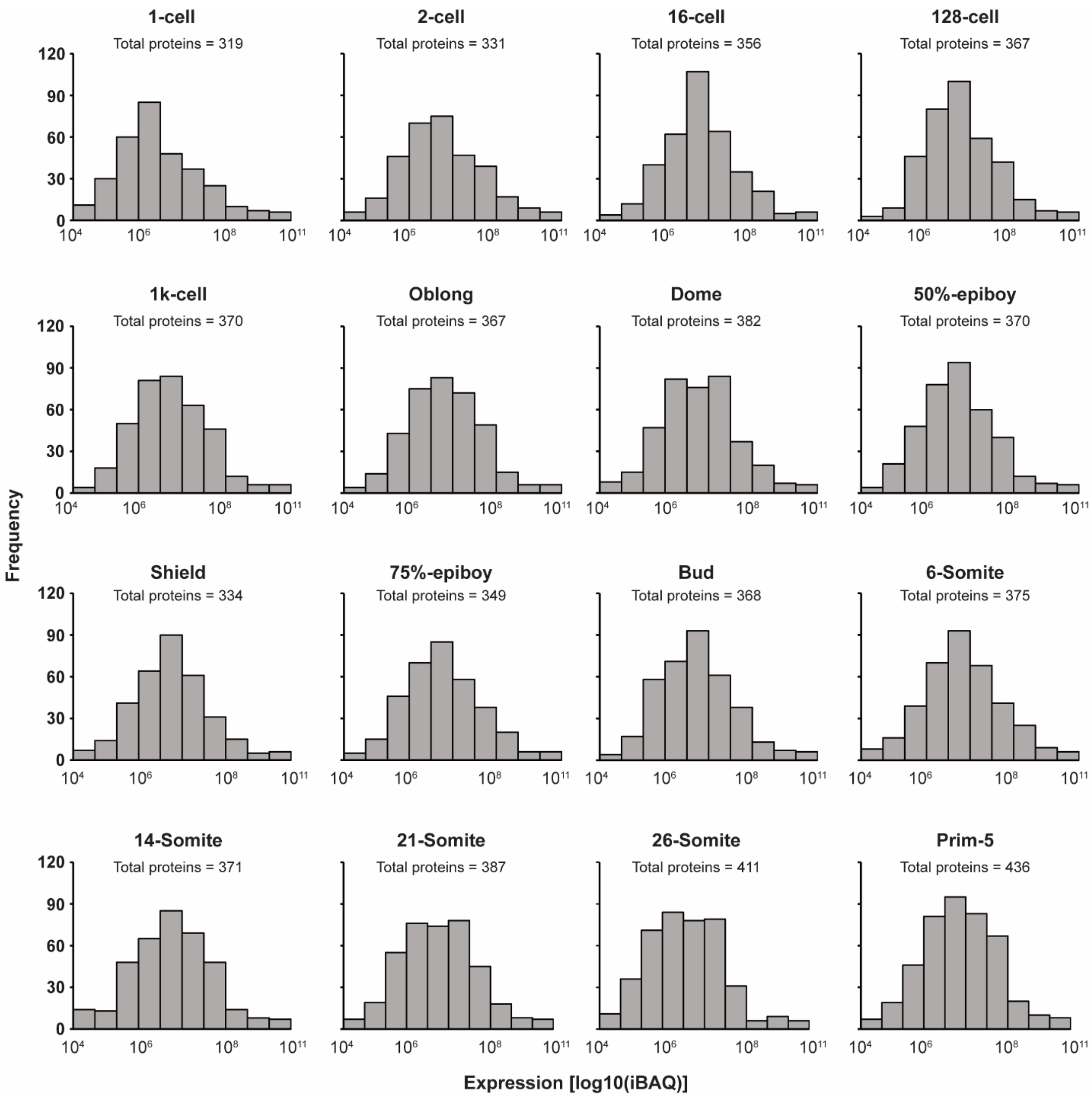


**Figure S2**. Histograms of the distribution of protein expression levels [log10(iBAQ)] at each stage. The total number of genes represented for the stage is shown on each panel. In the early stages, the distribution is unimodal and shifts toward higher abundance proteins and becomes more evenly distributed as development proceeds.

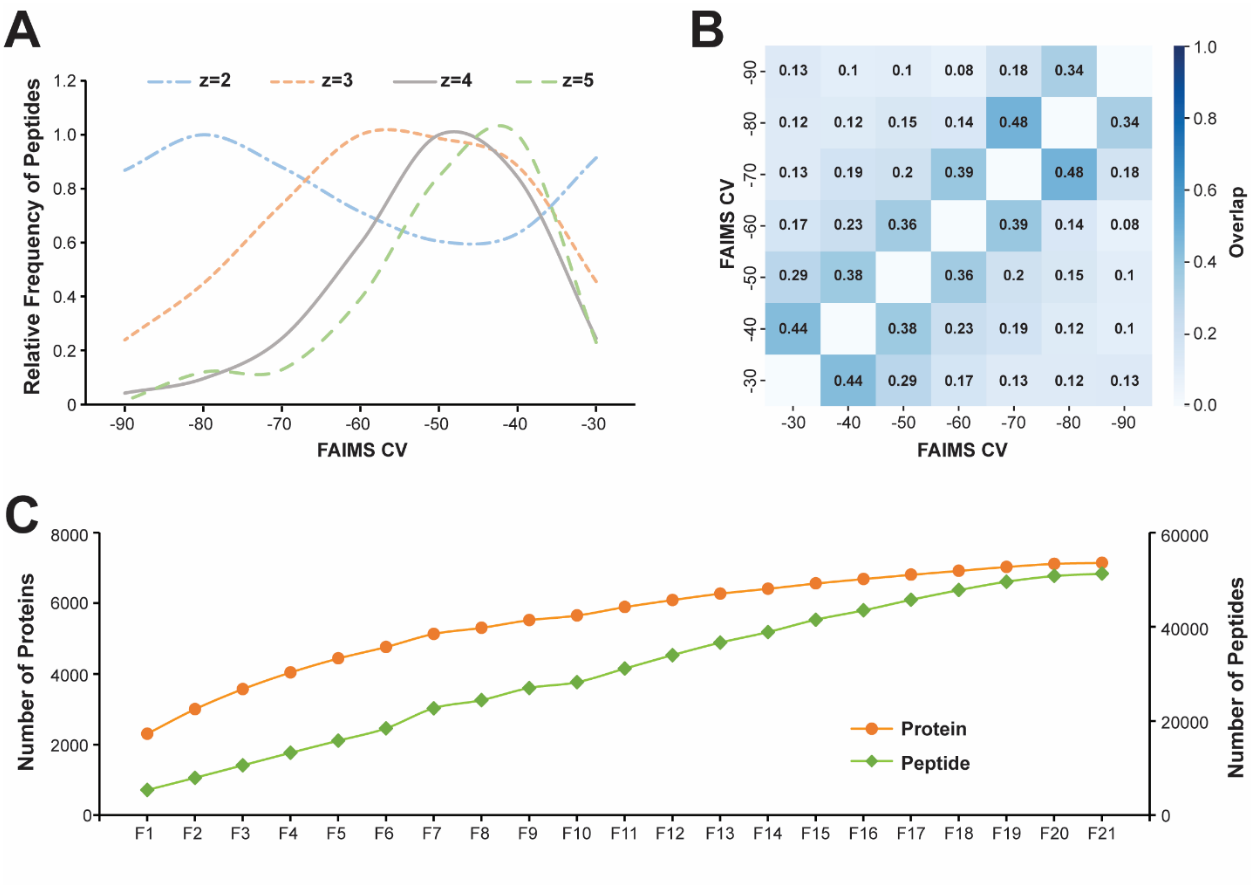


**Figure S3**. Peptide and protein identification from different CVs and high-pH RPLC fractions. The data is from Proteome Discoverer. (A) Distribution of the charge of the peptides across CV settings from -30 to -90 V. (B) Heat map of the percentage overlap of the peptides across CV settings from -30 to -90 V. (C) The accumulation number of peptide and protein identifications obtained from 21 high-pH RPLC fractions.

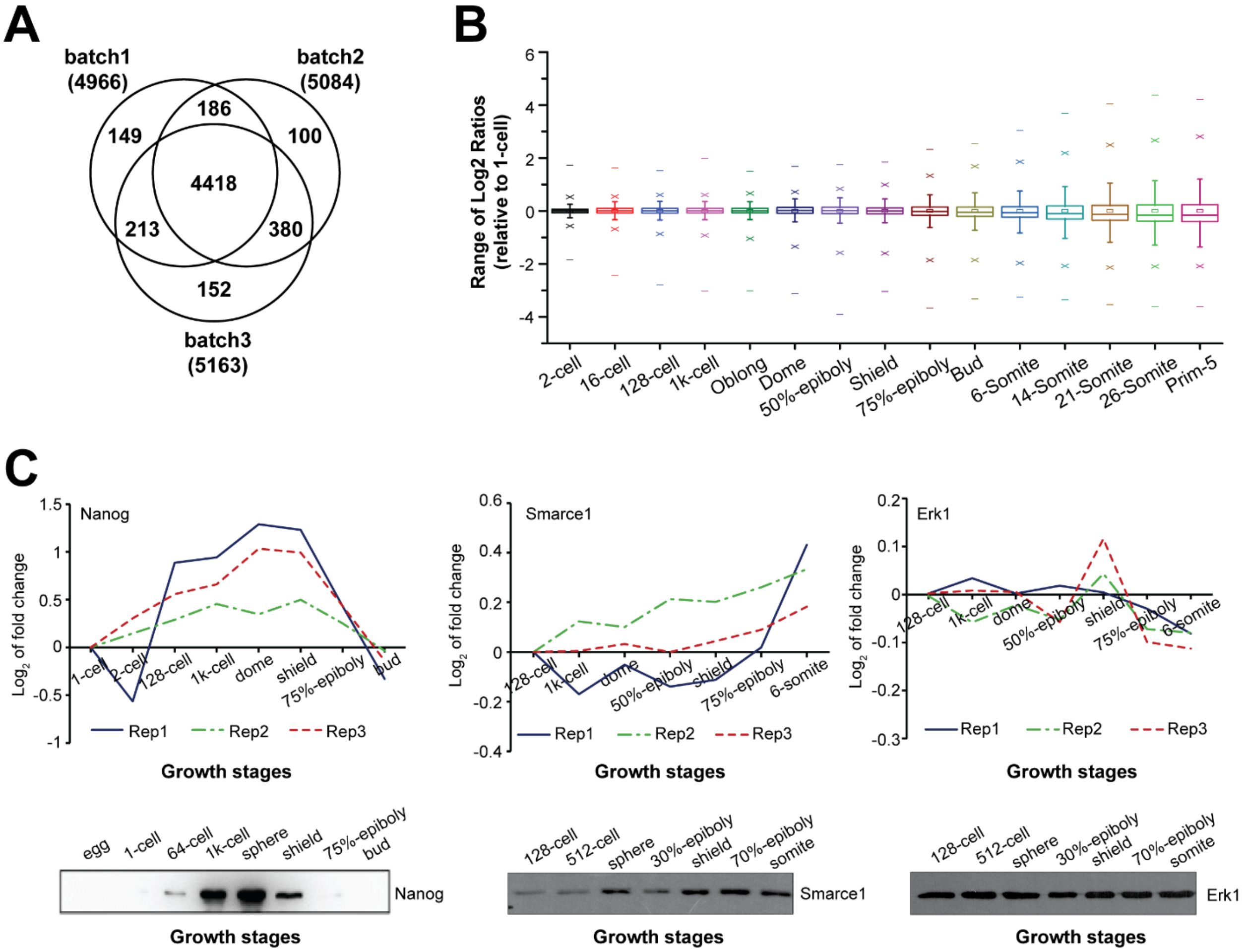


**Figure S4.** (A) Overlaps of proteins quantified from three biological replicates with MaxQuant software. (B) Boxplots of $\log_2$(reporter ion intensity ratios) at the 2-cell to Prim-5 stages. The reporter ion intensities of those 15 stages were normalized to the 1-cell stage. (C) Protein profiles of Nanog, Smarce1, and Erk1 obtained from this study correlated well with recently published Western blot data[1,2].

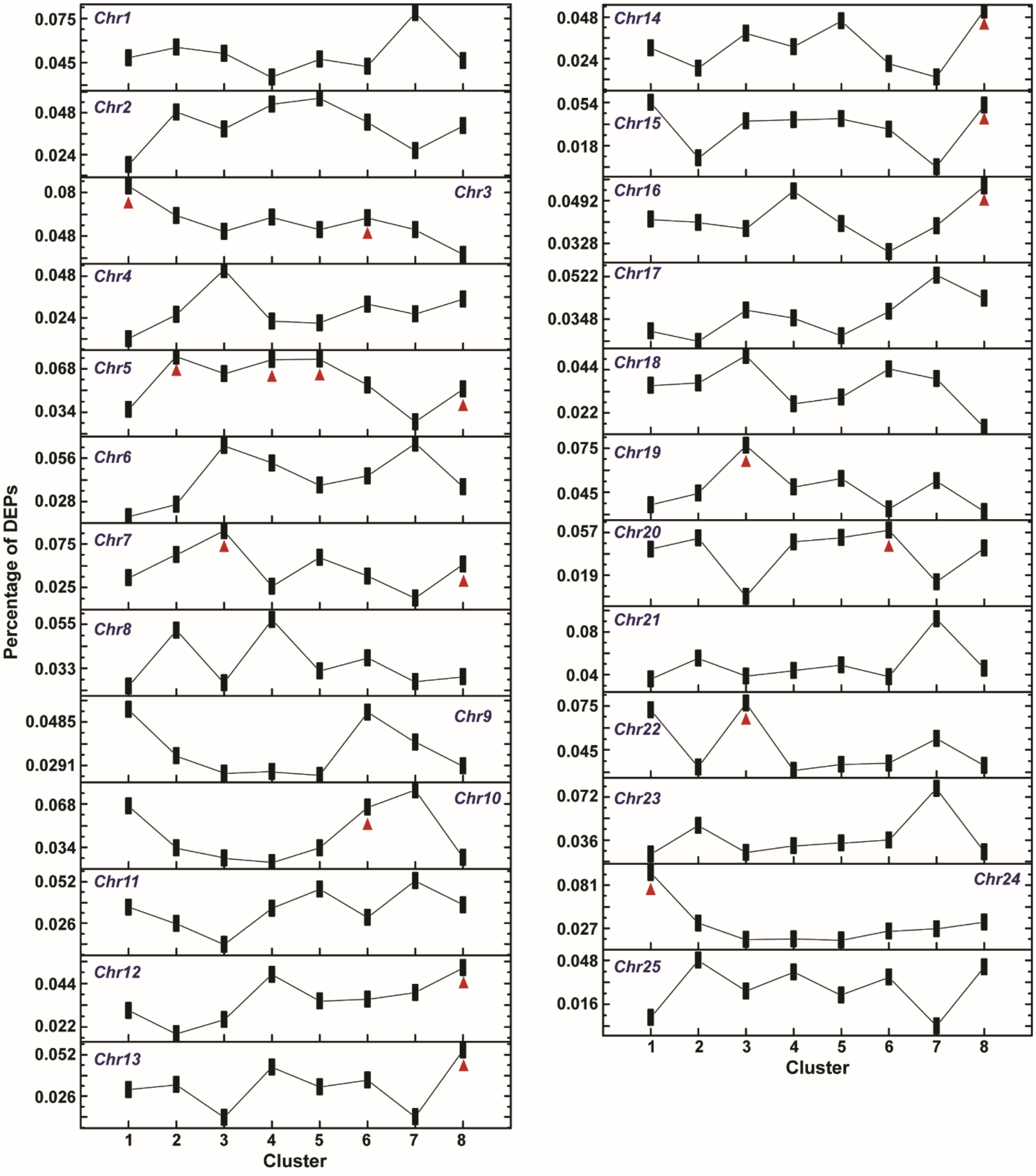


**Figure S5.** The distribution of differentially expressed proteins encoded by genes on each chromosome. The red arrows indicate the clusters showing a significant portion on each chromosome.

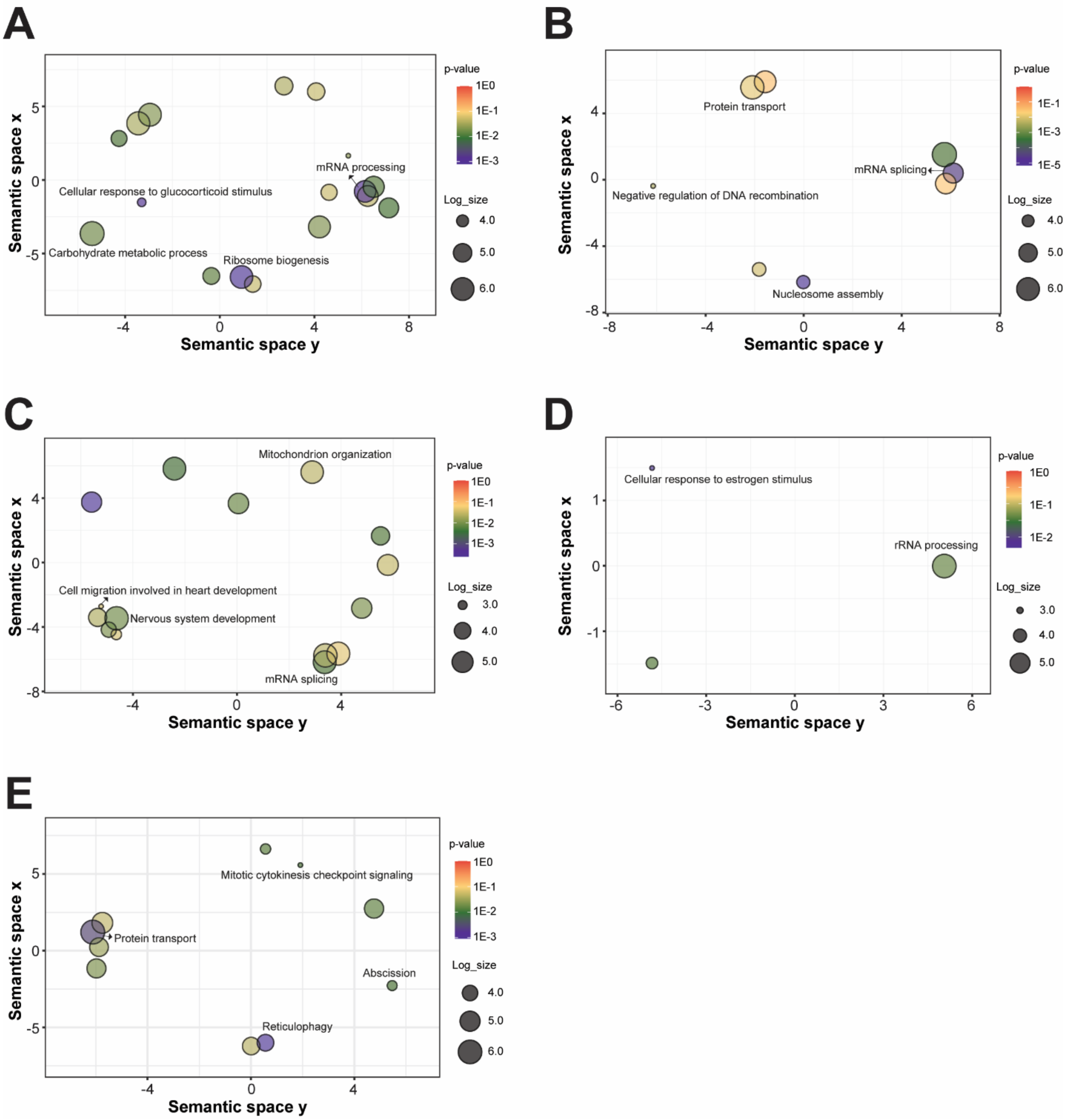


**Figure S6.** GO enrichment analysis for the DEPs located on (A) chromosome 5, (B) chromosome 7, (C) chromosome 3, (D) chromosome 22, and (E) chromosome 24. The REVIGO graph[3] displays the biological process enriched with DAVID Bioinformatics Resources. Each sphere represents a GO term colored by the GOfuncR enrichment in the –log10(*P*-value) scale. The semantic similarity of each GO term is represented by the position of each sphere on the graph. Plot size (sphere size) indicates the frequency of each GO term in which it is found in the GO database (i.e., the larger the sphere is, the more general the term is).

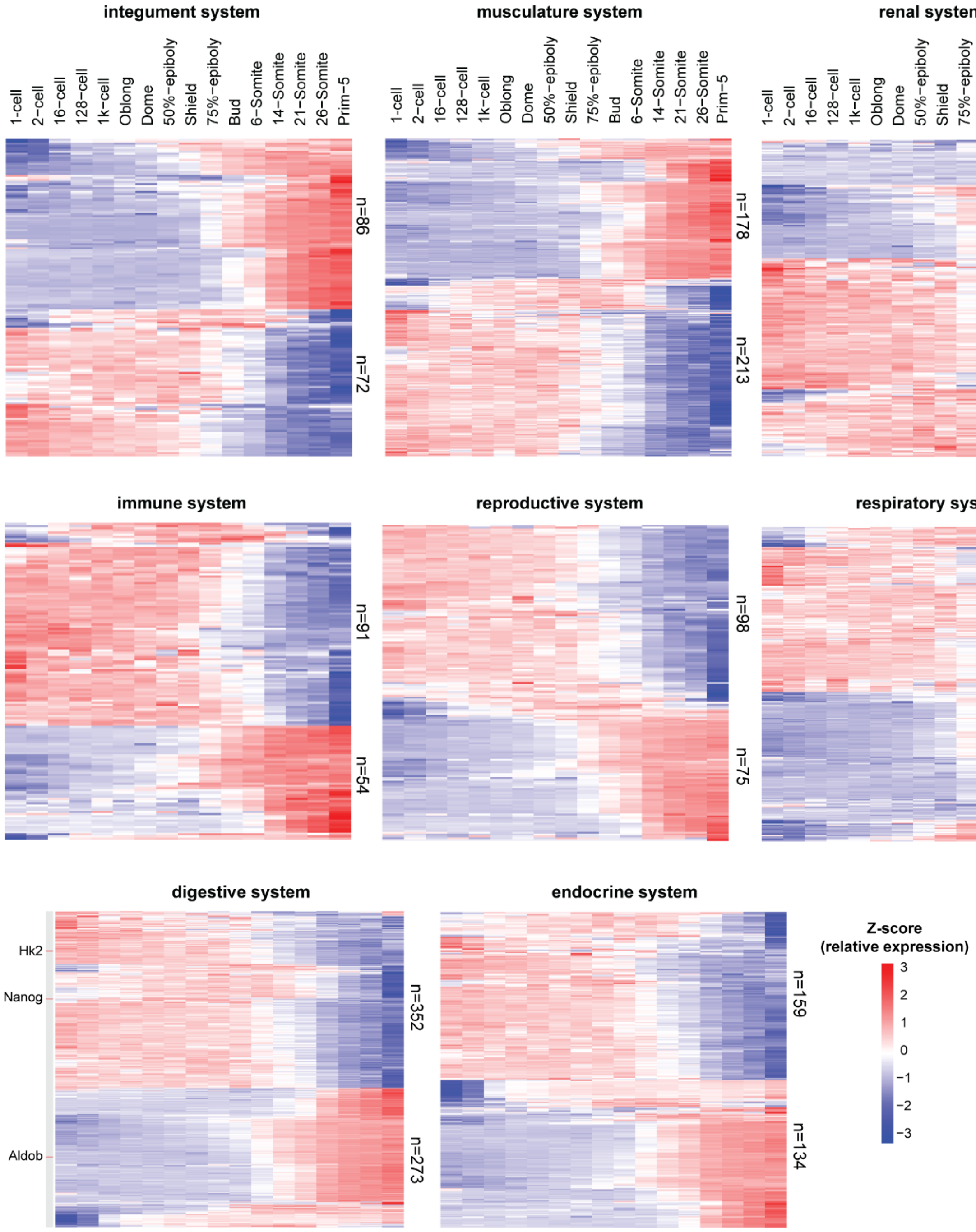


**Figure S7.** Heatmap of the expression profiles across the time course for differentially expressed proteins (DEPs) involved in different tissue systems. The numbers on the right represent the number of DEPs with an increased or decreased expression trend across embryogenesis.

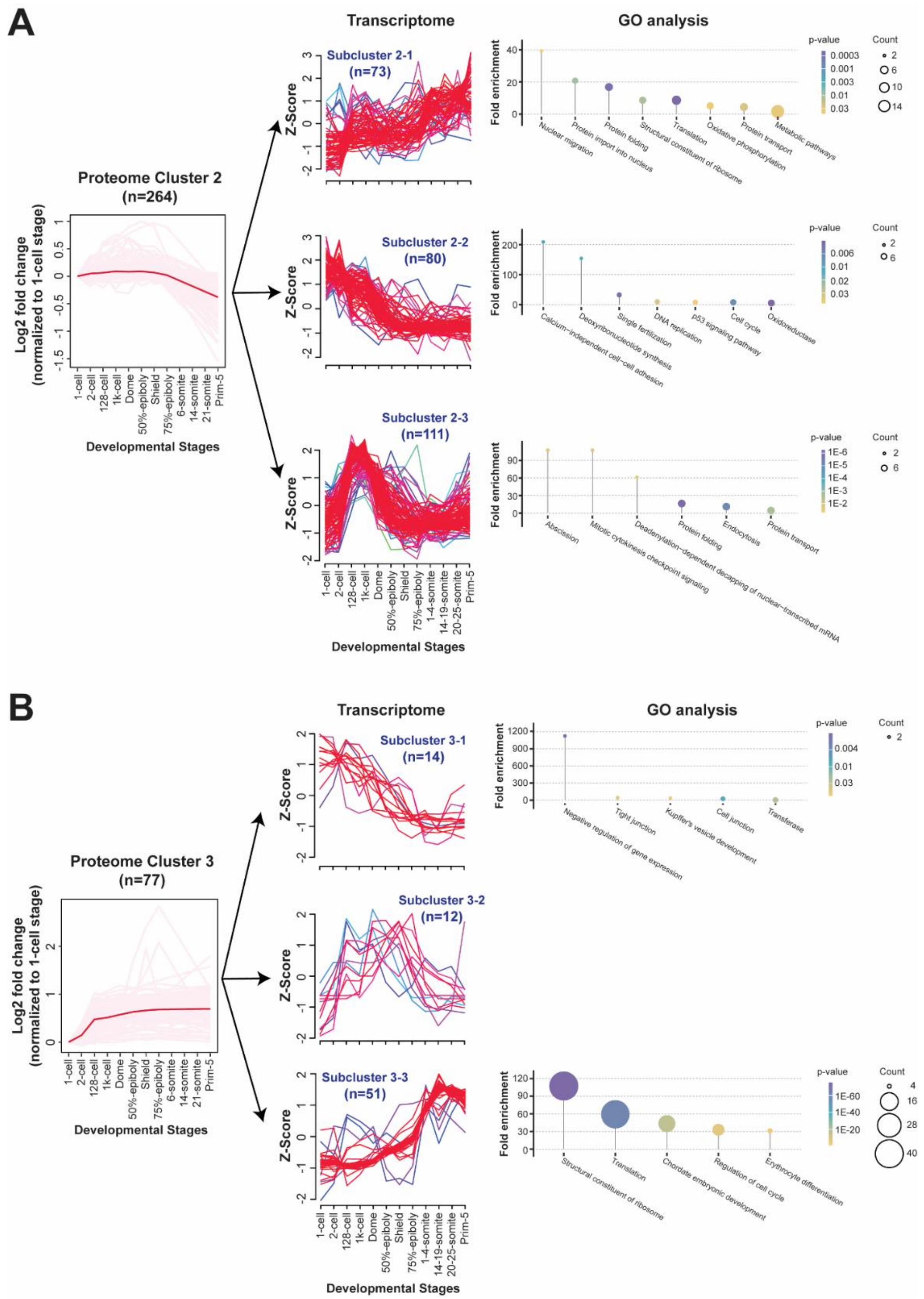


**Figure S8.** Proteome and transcriptome correlation data for the differentially expressed proteins (DEPs) in clusters 2 (A) and 3 (B). The genes of DEPs in clusters 2 and 3 are divided into three subclusters based on the dynamics of mRNA abundance across developmental stages. The twelve time points covered by the proteome and transcriptome data are the same as those shown in Figure 6. Gene ontology (GO) analysis was performed for genes in each subcluster, and the enriched biological processes are shown. Adjusted p-values and gene counts are also labeled.

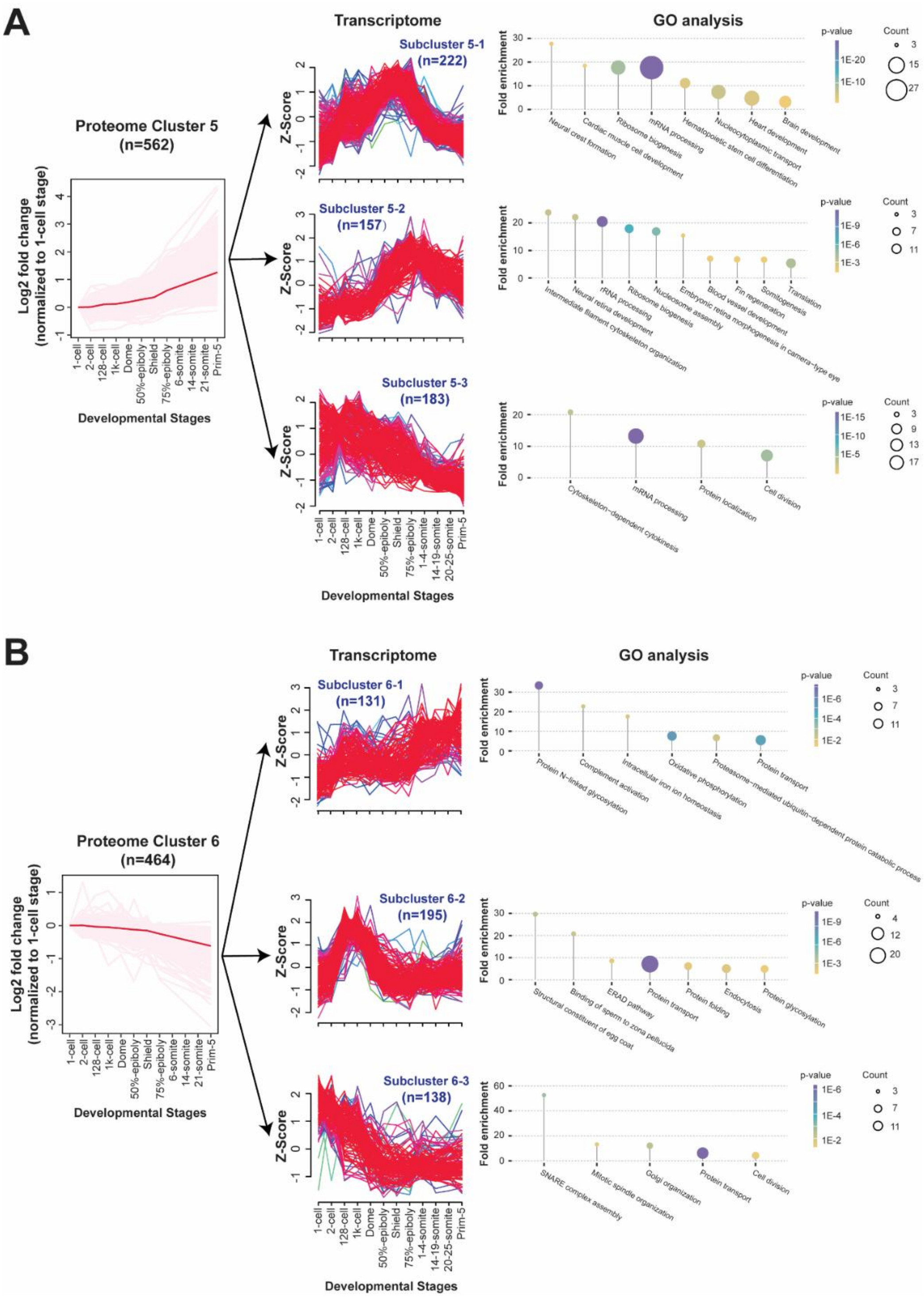


**Figure S9.** Proteome and transcriptome correlation data for the differentially expressed proteins (DEPs) in clusters 5 (A) and 6 (B). The genes of DEPs in clusters 5 and 6 are divided into three subclusters based on the dynamics of mRNA abundance across developmental stages. The twelve time points covered by the proteome and transcriptome data are the same as those shown in Figure 6. Gene ontology (GO) analysis was performed for genes in each subcluster, and the enriched biological processes are shown. Adjusted p-values and gene counts are also labeled.

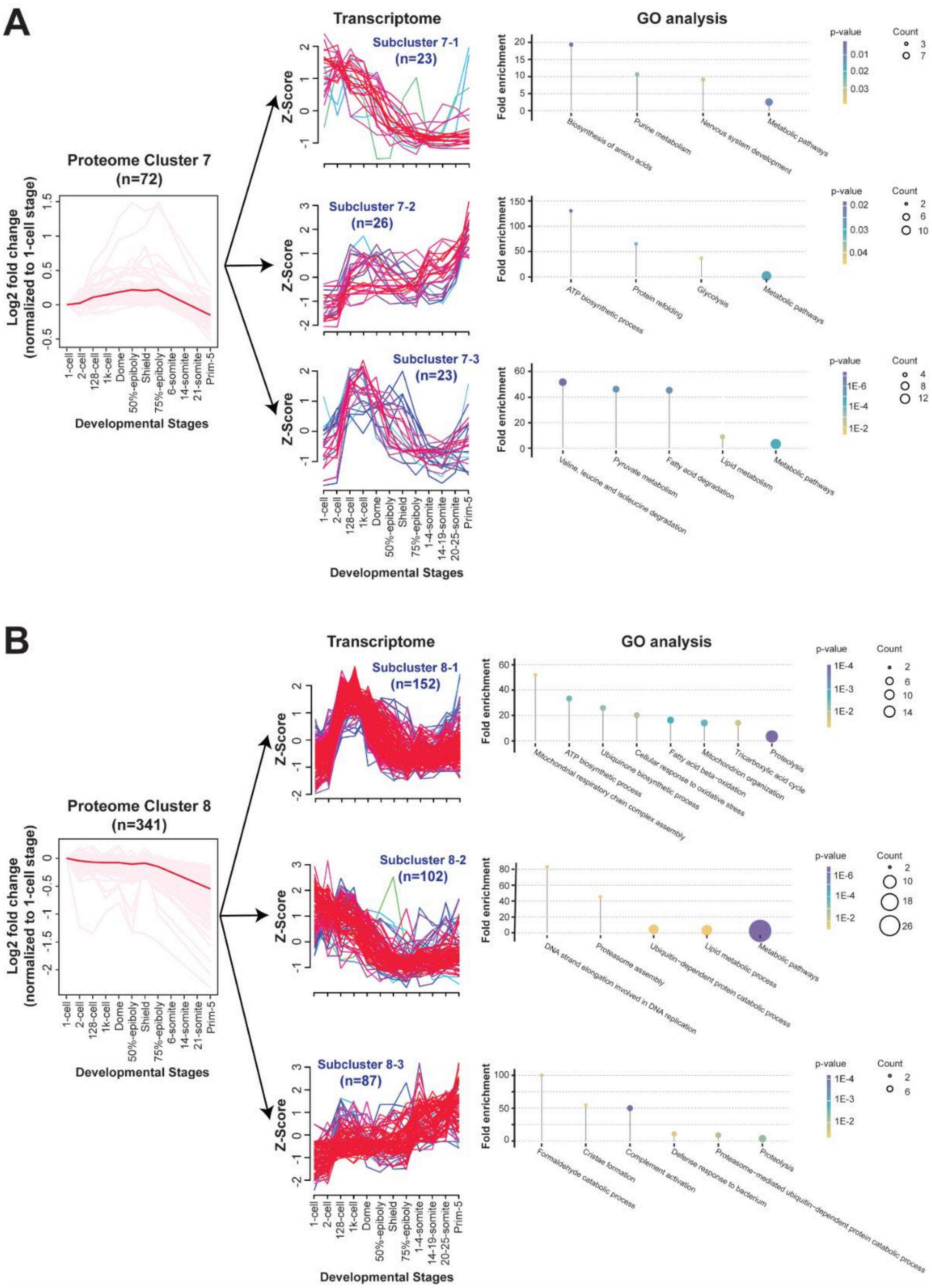


**Figure S10.** Proteome and transcriptome correlation data for the differentially expressed proteins (DEPs) in clusters 7 (A) and 8 (B). The genes of DEPs in clusters 7 and 8 are divided into three subclusters based on the dynamics of mRNA abundance across developmental stages. The twelve time points covered by the proteome and transcriptome data are the same as those shown in Figure 6. Gene ontology (GO) analysis was performed for genes in each subcluster, and the enriched biological processes are shown. Adjusted p-values and gene counts are also labeled.

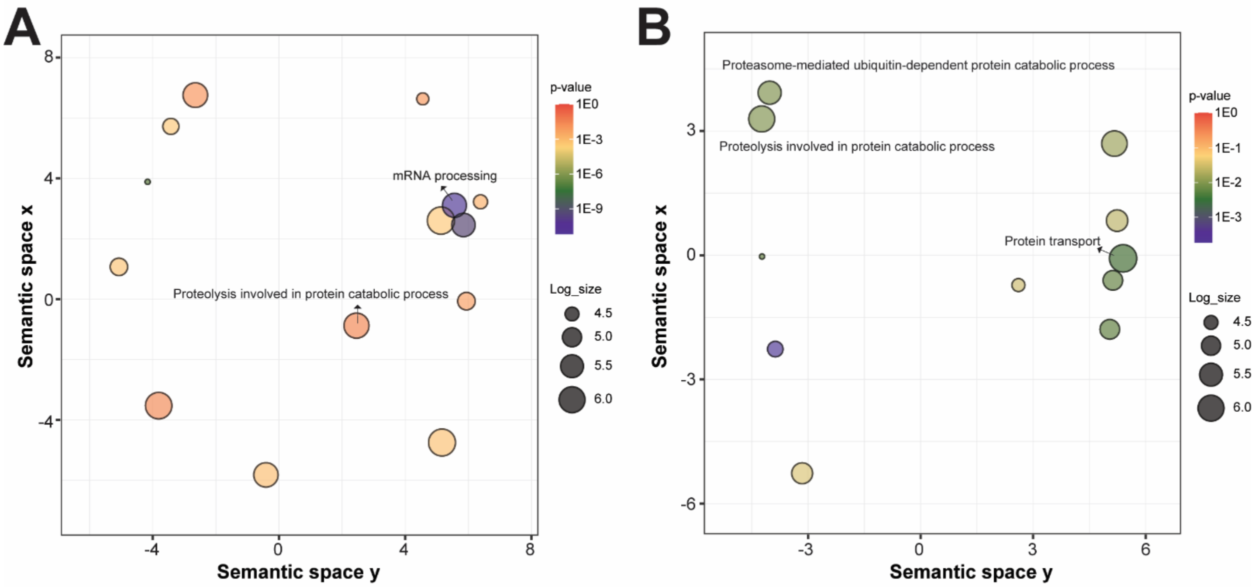


**Figure S11.** GO enrichment analysis for the DEPs in subcluster 5-3 (A) and subcluster 6-1 (B) with a strong negative correlation between mRNA and protein level. The REVIGO graph displays the biological process of GO enrichment. Each sphere represents a GO term colored by the GOfuncR enrichment in the –log10(*P*-value) scale. The semantic similarity of each GO term is represented by the position of each sphere on the graph. Plot size (sphere size) indicates the frequency of each GO term in which it is found in the GO database (i.e., the larger the sphere is, the more general the term is).

**Table S1.** The molecule types are revealed by the Ingenuity Pathway Analysis software.

| Type(s) | cluster1 | cluster2 | cluster3 | cluster4 | cluster5 | cluster6 | cluster7 | cluster8 |
|---|---|---|---|---|---|---|---|---|
| **Transcription regulator** | 4 | 13 | 6 | 9 | 64 | 11 | 2 | 9 |
| **Translation regulator** | 3 | 4 | 4 | 4 | 19 | 1 | 2 | 2 |
| **Transmembrane receptor** | 0 | 0 | 0 | 1 | 3 | 1 | 0 | 1 |
| **Transporter** | 2 | 21 | 1 | 14 | 18 | 37 | 4 | 21 |
| **Enzyme** | 26 | 75 | 10 | 56 | 90 | 111 | 37 | 118 |
| **Ion channel** | 2 | 1 | 0 | 2 | 0 | 3 | 0 | 3 |
| **Kinase** | 2 | 4 | 3 | 11 | 13 | 10 | 3 | 10 |
| **Peptidase** | 6 | 6 | 1 | 6 | 7 | 21 | 3 | 23 |
| **Phosphatase** | 2 | 2 | 0 | 3 | 7 | 6 | 0 | 3 |
| **Other** | 38 | 86 | 42 | 72 | 212 | 167 | 11 | 84 |

**Table S2.** The cellular component of proteins in each cluster was revealed by UniProt.

| Cellular component | cluster1 | cluster2 | cluster3 | cluster4 | cluster5 | cluster6 | cluster7 | cluster8 |
|---|---|---|---|---|---|---|---|---|
| **Mitochondrion** | 12 | 42 | 3 | 14 | 14 | 31 | 20 | 72 |
| **Nucleus** | 21 | 56 | 21 | 68 | 314 | 77 | 18 | 53 |
| **Cytoskeleton** | 2 | 5 | 1 | 11 | 28 | 8 | 2 | 6 |
| **ER/Golgi** | 15 | 29 | 1 | 17 | 11 | 100 | 3 | 29 |
| **Endosome/lysosome** | 6 | 16 | 0 | 7 | 3 | 32 | 0 | 19 |
| **Cytosol** | 19 | 40 | 5 | 33 | 39 | 45 | 12 | 32 |

### Reference


1. Schnabel, D., Castillo-Robles, J., and Lomeli, H. (2019). Protein Purification and Western Blot Detection from Single Zebrafish Embryo. Zebrafish *16*, 505–507. https://doi.org/10.1089/zeb.2019.1761.

2. He, M., Zhang, R., Jiao, S., Zhang, F., Ye, D., Wang, H., and Sun, Y. (2020). Nanog safeguards early embryogenesis against global activation of maternal β-catenin activity by interfering with TCF factors. PLoS Biol *18*, e3000561. https://doi.org/10.1371/journal.pbio.3000561.

3. Supek, F., Bošnjak, M., Škunca, N., and Šmuc, T. (2011). REVIGO summarizes and visualizes long lists of gene ontology terms. PLoS One *6*, e21800. https://doi.org/10.1371/journal.pone.0021800.
